# Activation of NIX-mediated mitophagy and replication by an interferon regulatory factor homologue of human herpesvirus

**DOI:** 10.1101/506113

**Authors:** Mai Tram Vo, Barbara J Smith, John Nicholas, Young Bong Choi

**Affiliations:** Department of Oncology, Sidney Kimmel Comprehensive Cancer Center, Johns Hopkins University School of Medicine, Baltimore, Maryland, United States of America; Department of Cell Biology, Institute for Basic Biomedical Sciences, Johns Hopkins University School of Medicine, Baltimore, Maryland, United States of America

## Abstract

Viral control of mitochondrial quality and content has emerged as an important mechanism for counteracting the host response to virus infection. Despite the knowledge of this crucial function of some viruses, little is known about how herpesviruses regulate mitochondrial homeostasis during infection. Human herpesvirus 8 (HHV-8) is an oncogenic virus causally related to AIDS-associated malignancies. Here, we found that HHV-8-encoded viral interferon regulatory factor 1 (vIRF-1) promotes mitochondrial clearance by activating mitophagy to support virus replication. Genetic interference with vIRF-1 expression or targeting to the mitochondria inhibits HHV-8 replication-induced mitophagy and leads to an accumulation of mitochondria. Moreover, vIRF-1 binds directly to a mitophagy receptor, NIX, on the mitochondria and activates NIX-mediated mitophagy to promote mitochondrial clearance. Genetic and pharmacological interruption of vIRF-1/NIX-activated mitophagy inhibits HHV-8 productive replication. In short, our findings uncover an essential role of vIRF-1 in mitophagy activation for successful lytic replication of HHV-8.

## Introduction

Mitochondria are vital energy-generating organelles in eukaryotic cells. They also integrate signals controlling multiple cellular functions such as apoptosis, cell cycle, and development. In addition, recent studies have shown that mitochondria serve as a platform for mediating antiviral signaling pathways that lead to apoptosis and innate immune responses to virus infection^1,2^. For example, proapoptotic BH3-only proteins (BOPs) are elevated^3^ and/or activated during virus replication and induce mitochondrial outer membrane permeabilization, a crucial step in the intrinsic apoptotic process that triggers the release of soluble apoptogenic factors from the intermembrane space. In the innate immune response to virus infection, the RIG-I-like receptors (RLRs) RIG-I and MDA-5 recognize cytosolic viral RNA and promote the oligomerization of the mitochondrial antiviral signaling adaptor protein (MAVS; also known as IPS-1, VISA, and Cardif)^4^, which recruits TBK1 and IKKi kinases to activate IRF3 and IRF7 transcription factors^2^. These activated IRFs induce the expression of type I interferon (IFN) genes, the products of which restrict virus replication. Therefore, successful virus infection and replication are in large part achieved by the ability of viruses to attenuate the innate antiviral responses mediated by mitochondria.

Particular viral proteins have evolved to enable virus infection and replication by modulating the mitochondria-mediated antiviral responses. For example, human herpesviruses encode anti-apoptotic proteins that inhibit the intrinsic apoptosis pathway^1^, and hepatitis C virus encodes a serine protease NS3/NS4A that disrupts RLR signaling and IFN-β production by cleaving MAVS^5^. On the other hand, mitochondria generate reactive oxygen species (ROS) as a byproduct of electron transport complexes. Accumulation of damaged or dysfunctional mitochondria leads to aberrant ROS generation, which can exacerbate apoptosis and augment RLR-MAVS signaling^6,7^. Several viruses are known to effect elimination of infection-altered mitochondria via mitophagy, potentially attenuating apoptosis and innate immune responses^8^.

Mitophagy is a key mechanism of mitochondrial quality control that eliminates aged, dysfunctional, damaged or excessive mitochondria via selective autophagy^9,10^. A recent study indicates that impaired removal of damaged mitochondria by mitophagy can lead to STING-dependent inflammation and neurodegeneration^11^. PINK1/PARK2-mediated mitophagy is one of the most well-characterized; after the loss of mitochondrial membrane potential, PINK1 is stabilized on the outer mitochondrial membrane (OMM), where it phosphorylates both ubiquitin (Ub) and the Ub-like domain of PARK2, leading to its activation^12-14^. In turn, activated PARK2 ubiquitinates mitochondrial proteins such as MFN1/2, DRP1, BCL-2, and VDAC1. Mitophagy receptors such as NDP52 and OPTN bind to and link the ubiquitinated mitochondria to autophagosomes via interaction with autophagy-related protein 8 (ATG8) family members such as LC3 and GABARAP^15^. PARK2-independent mitophagy pathways mediated by other mitophagy receptors such as NIX/BNIP3L, BNIP3, FUNDC1, and PHB-2 have been identified under certain physiological conditions^16-19^. Nonetheless, viral regulation of mitophagy is an understudied area of investigation.

Human herpesvirus 8 (HHV-8) is a pathogen associated with Kaposi’s sarcoma (KS), an endothelial cell disease, the B cell malignancy primary effusion lymphoma (PEL) and the lymphoproliferative disease multicentric Castleman’s disease^20,21^. HHV-8 productive replication, in addition to latency, is important for maintaining viral load within the host and for KS pathogenesis. Successful HHV-8 replication is in large part achieved by the ability of the virus to inhibit apoptosis and innate immune responses elicited by infection of host cells. HHV-8 encodes a number of proteins expressed during the lytic cycle that have demonstrated or potential abilities to promote virus productive replication via inhibition of antiviral responses activated by infection-induced stress. Amongst them, viral IFN regulatory factor 1 (vIRF-1) is believed to play important roles in blocking IFN and other stress responses to virus infection and replication through inhibitory interactions with cellular stress signaling proteins such as p53, ATM, IRF1, GRIM19, and SMAD3/4^22,23^. Furthermore, we found that vIRF-1 localizes to mitochondria upon virus replication and suppresses mitochondria-mediated apoptosis and innate immune responses via its inhibitory interactions with proapoptotic BOPs and MAVS^3,24,25^. However, the precise role of mitochondria-localized vIRF-1 in HHV-8 biology remains elusive. In this study, we identify a new function of vIRF-1 in its activation of mitophagy and support of virus replication via this pathway.

## Results

### vIRF-1 downregulates mitochondrial content during lytic reactivation

We previously reported that the level of MAVS is significantly diminished in the HHV-8-infected PEL cell line, BCBL-1, that expresses vIRF-1 following lytic reactivation^25^. Another mitochondrial protein TOM20 was also strongly reduced in the mitochondria of vIRF-1-expressing lytic cells compared to that in vIRF-1-negative neighbor cells (Supplementary Fig. 1a). Moreover, MitoTracker Red staining of mitochondria showed an apparent decrease in the levels of mitochondria of vIRF-1-expressing lytic cells (Supplementary Fig. 1b), implying that the reduced levels of MAVS and TOM20 might result from a loss of mitochondrial content rather than a specific inhibition of the expression or stability of the proteins. To further address this notion, we examined the levels of mitochondrial DNA (mtDNA) using a combined approach of immunofluorescence assay (IFA) and fluorescent *in situ* hybridization (FISH). Indeed, the result showed that mtDNA was less readily detected in vIRF-1-expressing lytic cells (Fig. 1a). Furthermore, immunoblotting and image cytometry analyses showed that the expression of MTCO2 (mtDNA-encoded cytochrome c oxidase II) was significantly reduced in lytic TRExBCBL-1-RTA (hereafter simply termed iBCBL-1) cells, doxycycline (Dox)-inducible for lytic switch protein RTA^26^, that were treated with Dox for 2 days (Figs. 1b and 1c). These results suggest that mitochondria content of cells could be downregulated, conceivably by vIRF-1, following lytic reactivation of HHV-8.

**Fig. 1.**
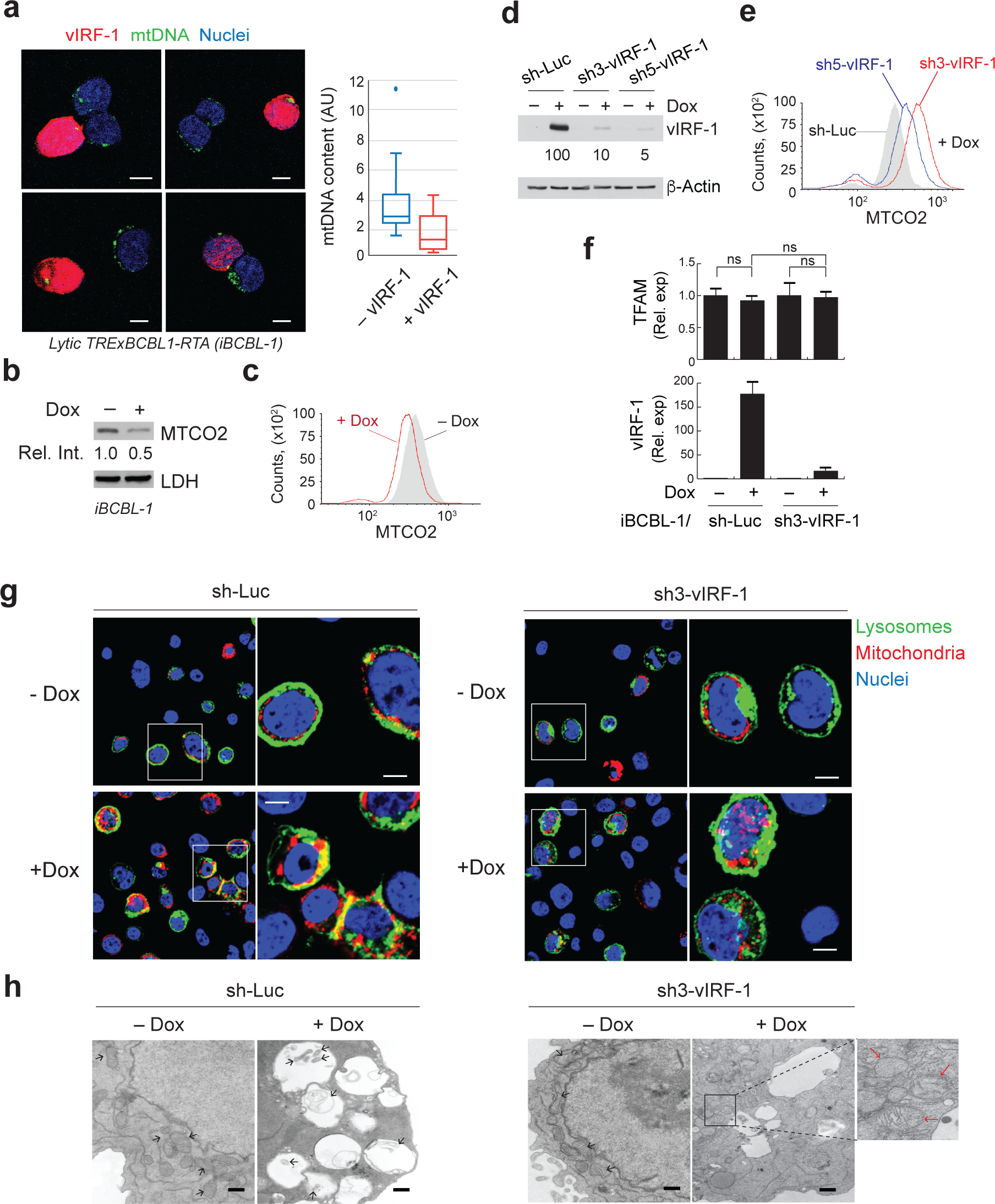
vIRF-1 downregulates mitochondria content via mitophagy during lytic replication. **a** Quantitative assessment of mitochondrial DNA (mtDNA) in lytic iBCBL-1 cells, which were treated with 1 µg/ml Dox for 2 days; a combined approach of immunofluorescent assay (for detection of vIRF-1 protein) and fluorescent *in situ* hybridization (for detection of mtDNA) was used. Sixteen images were taken randomly, and representative images are presented. The levels of mtDNA in vIRF-1-negative (– vIRF-1) and -positive (+ vIRF-1) cells were determined using ImageJ software and are shown in a Box-Whisker diagram; AU, arbitrary units. Scale bar, 5 µm. **b,c** Immunoblotting and image cytometric analyses of MTCO2 expression in iBCBL-1 cells that were left untreated or treated with Dox for 2 days. Lactate dehydrogenase (LDH) was used as loading control. **d** Validation of Dox-inducible expression of vIRF-1 shRNA3 (sh3-vIRF-1) and shRNA5 (sh5-vIRF-1) along with control luciferase shRNA (sh-Luc) in iBCBL-1 cells using immunoblotting. β-Actin was used as loading control. The relative intensities of vIRF-1 normalized to β-Actin were noted below the vIRF-1 blot panel. **e** Image cytometric analysis of MTCO2 expression in control and vIRF-1-depleted iBCBL-1 cells that were treated with Dox for 2 days. **f** Real-time quantitative PCR (RT-qPCR) analysis of the mRNA expression of TFAM and vIRF-1 in control and vIRF-1-depleted iBCBL-1 cells that were left untreated or treated with Dox for 2 days. The relative expression levels (Rel. exp) were calculated using comparative Ct (2^-ΔΔCt^) method and normalized to 18S rRNA levels. **g** Assessment of the formation of mitochondria-containing lysosomes (mitolysosomes) in control and vIRF-1-depleted iBCBL-1 cells, which were left untreated or treated with Dox for 2 days, using CellLight^®^ BacMam 2.0 system (see STAR Methods for details). Representative images of each sample are presented, and the yellow areas indicate mitolysosomes. Scale bar, 5 µm. **h** Electron microscopy images of control and vIRF-1-depleted iBCBL-1 cells that were left untreated or treated with Dox for 2 days. Black and red arrows indicate mitochondria with normal and disrupted cristae, respectively. Scale bar, 500 nm.

To next examine whether vIRF-1 is indeed involved in the regulation of mitochondria content, we generated a cell line that stably expresses a short hairpin RNA (shRNA) directed against vIRF-1 in iBCBL-1 cells along with a control cell line expressing shRNA directed against luciferase (Luc). Constitutive knockdown of vIRF-1 was previously shown to induce death of latently HHV-8-infected PEL cell lines^27^. To circumvent this limitation, we used an inducible system; Dox-inducible vIRF-1 shRNAs, sh3 and sh5, resulted in 10-to 20-fold decreases in vIRF-1 protein expression compared to control shRNA (sh-Luc) in lytic iBCBL-1 cells (Fig. 1d). We next assessed the mitochondria content of lytic control and vIRF-1-depleted iBCBL-1 cells. Image cytometry analysis showed that, when vIRF-1 was depleted, lytic cells with higher levels of MTCO2 were increased in number (Fig. 1e). Taken together, the results suggest that vIRF-1 plays an important role in the downregulation of mitochondria content during virus replication.

### vIRF-1 controls mitochondria content via mitophagy

The cellular content of mitochondria is fundamentally controlled by the balance of anabolic biogenesis and catabolic degradation such as selective autophagy of mitochondria (termed mitophagy). Mitochondrial biogenesis is a complex process requiring the coordinated expression of nuclear and mitochondrial DNAs. Nucleus-encoded mitochondrial transcription factor A (TFAM) is essential for transcription and replication of mtDNA, which are important steps in the mitochondrial biogenesis^28^. Thus, we examined the transcriptional levels of TFAM but observed no significant difference between control and vIRF-1-depleted iBCBL-1 cells that were left untreated or treated with Dox (Fig. 1f), suggesting that virus replication and vIRF-1 might not influence mitochondrial biogenesis. Therefore, we extrapolated that mitophagy may be involved in vIRF-1 regulation of mitochondrial content during lytic replication. To test this notion, we first examined whether HHV-8 activates mitophagy following lytic reactivation. The results showed that autophagy inhibitors including bafilomycin A1 (Baf A1) and leupeptin blocked a decrease in the MTCO2 levels in Dox-treated (lytically-reactivated) iBCBL-1 cultures (Supplementary Figs. 1c and 1d). Consistent with this, immunoblotting analysis showed that a decrease of MTCO2 protein was inhibited by Baf A1 and chloroquine (CQ), another autophagy inhibitor, but not by a proteasome inhibitor, MG132, in Dox-induced iBCBL-1 cultures (Supplementary Fig. 1e). We further examined the formation of mitochondria-containing autolysosomes (hereafter simply named mitolysosomes), an end-point readout of mitophagy^29^, using CellLight^TM^ BacMam-labeling of mitochondria and lysosomes (see Material and Methods for details). The results showed that the presence of mitolysosomes was more evident in lytic control iBCBL-1 cells than latent control and both latent and lytic vIRF-1-depleted iBCBL-1 cells (Fig. 1g). Furthermore, electron microscopy imaging demonstrated the apparent presence of mitolysosomes in lytic control cells but not in lytic vIRF-1-depleted iBCBL-1 cells (Fig. 1h). It is noteworthy that mitochondria with disrupted cristae were often observed outside the autophagic vacuoles of lytic vIRF-1-depleted iBCBL-1 cells (Fig. 1h, red arrows). Taken together, our results suggest that vIRF-1 is likely to be involved in activation of mitophagy to control mitochondrial content of cells during virus replication.

### vIRF-1 activates NIX-mediated mitophagy

Mitophagy is triggered by activation of specific autophagy receptors localized mainly on the outer mitochondrial membrane (OMM); these proteins interact with ATG8 family members including LC3 and GABARAP via a short-linear motif termed the ATG8-interacting motif (AIM) or LC3-interacting region (LIR), which forms a bridge linking the mitochondria to the autophagosomes^10^. Thus, we hypothesized that vIRF-1 may promote mitophagy by recruiting the mitophagy machinery and/or activating them on the mitochondria. Firstly, we investigated changes in the levels of the mitophagy machinery including mitophagy receptors and LC3 on the mitochondria isolated from latent and lytic iBCBL-1 cells. Consistent with our previous report^25^, vIRF-1 was readily detected in the mitochondrial fraction isolated from lytic iBCBL-1 cells (Fig. 2a). When autophagy is induced, LC3 is processed from a cytosolic form, LC3-I (18 kDa), to the LC3-II (16 kDa) form that is lipidated with phosphatidylethanolamine and associated with the autophagic vesicle membranes. Intriguingly, the LC3B-II form, but not the LC3B-I form, was readily detected in the mitochondria and here exhibited a more than 2-fold increase upon virus replication while total LC3B levels remained unchanged after lytic reactivation (Fig. 2a), indicating that mitophagy may be selectively induced during virus replication. We next examined the levels of cellular mitophagy receptors including NIX (also termed BNIP3L), OPTN, NDP52, p62, NBR1, and FUNDC1. Interestingly, the level of mitochondria-associated NIX was upregulated by more than 2-fold while the other receptors remained essentially unchanged (Fig. 2a). The mitochondrial fission protein DRP1 and the mitochondrial chaperone HSP60 were used as a loading control (Fig. 2a). NIX mRNA expression was not induced by lytic reactivation (Fig. 2b), indicating that NIX protein may be stabilized and/or translocated to the mitochondria during HHV-8 replication. We failed to detect expression of another mitophagy receptor BNIP3, a paralog of NIX, in both latent and lytic iBCBL-1 cells. At present we cannot rule out roles of other mitophagy receptors other than NIX in virus replication-induced mitophagy activation. Nonetheless, based on our finding of NIX regulation by lytic induction, we investigated further the function of NIX for virus replication- and vIRF-1-induced mitophagy activation in subsequent studies.

**Fig. 2.**
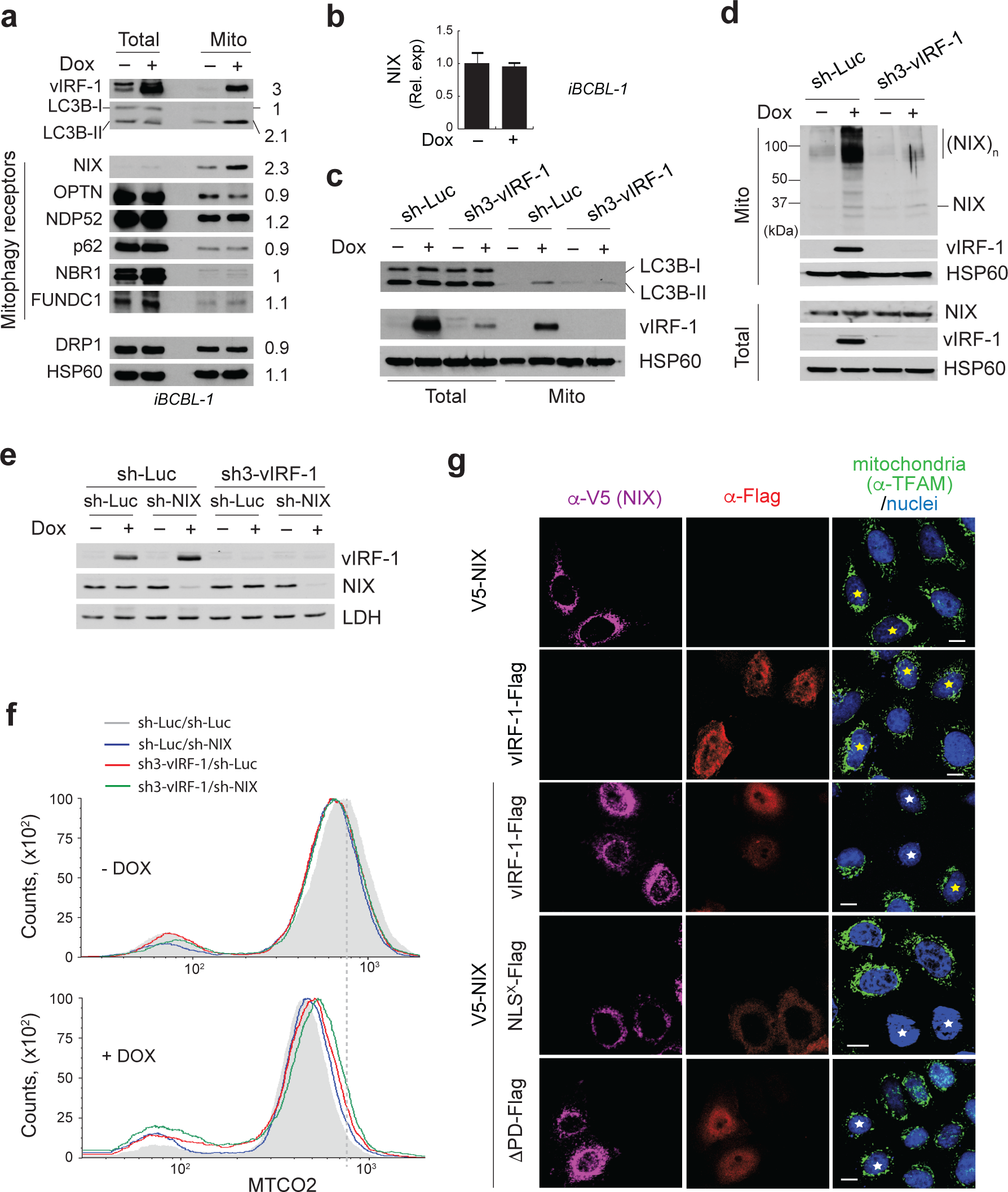
vIRF-1 activates NIX-mediated mitophagy. **a** Immunoblots of total-cell and mitochondrial extracts derived from iBCBL-1 cells that were left untreated or treated with Dox for 2 days. The relative band intensity of the Dox-treated mitochondrial fraction compared to the control mitochondrial fraction without Dox was calculated and shown in the column (right). HSP60 was used as loading control. **b** RT-qPCR analysis of the mRNA expression of NIX in latent and lytic iBCBL-1 cells. The relative expression levels (Rel. exp) were calculated using the comparative Ct method. **c, d** Immunoblots of total-cell and mitochondrial extracts derived from control and vIRF-1-depleted iBCBL-1 cells that were left untreated or treated with Dox for 2 days. (NIX)_n_ indicates a putative dimerized or polymerized form of NIX. **e** Validation of Dox-induced depletion of NIX and/or vIRF-1 in iBCBL-1 cells using immunoblotting. The cells were left untreated or treated with Dox for 2 days. LDH was used as loading control. **f** Image cytometric analysis of MTCO2 expression in latent and lytic iBCBL-1 cells that were depleted of NIX and/or vIRF-1. The cells were left untreated or treated with Dox for 4 days. The dotted line shows the peak for MTCO2 in latent cells (no Dox) doubly-transduced with sh-Luc. **g** IFA analysis of mitochondria content (TFAM) in HeLa.Kyoto cells that were transiently transfected for 24 h with V5-tagged NIX (V5-NIX) and/or Flag-tagged vIRF-1 (wild type and its derivatives, NLS^X^ and ΔPD, that are defective in the nuclear localization signal and the proline-rich domain (PD), respectively). Singly- and dually-transfected cells are marked with yellow and white stars, respectively. Scale bar, 10 µm.

We next examined whether vIRF-1 is required for the recruitment of LC3B-II and NIX to the mitochondria. Immunoblotting analyses showed that virus replication-induced translocation of LC3B-II to the mitochondria was abated when vIRF-1 was depleted (Fig. 2c). Similarly, virus replication-promoted mitochondrial localization of NIX and formation of its shifted bands to about 100-kDa, which may represent a homodimer of NIX, were significantly reduced in lytic vIRF-1-deficient iBCBL-1 cells (Fig. 2d). Whether NIX dimerization is required for mitophagy activation remains unclear; however, hypoxia-induced dimerization of a paralog of NIX, BNIP3, was proposed to be related to mitophagy activation *in vivo*^18^. Therefore, we hypothesized that vIRF-1 may activate NIX-mediated mitophagy via promotion of NIX recruitment to and dimerization on mitochondria. To test this possibility, we transiently transfected HeLa. Kyoto cells with Flag-tagged vIRF-1 (vIRF-1-Flag) and/or V5-tagged NIX (V5-NIX). Contrary to our expectation, there was no effect of vIRF-1 on promotion of V5-NIX recruitment to and dimerization on mitochondria (Supplementary Fig. 2). To examine whether NIX is involved functionally in vIRF-1 regulation of mitochondria content, we stably transduced control and vIRF-1-depleted iBCBL-1 cells with Dox-inducible NIX shRNA (sh-NIX). Dox-inducible depletion of endogenous NIX and vIRF-1 proteins were verified using immunoblotting in each cell line (Fig. 2e). Image cytometric analysis showed that depletion of NIX alone had a minor effect on the mitochondrial content, but double depletion of NIX and vIRF-1 significantly increased mitochondrial content in lytic cells to an even greater extent than depletion of vIRF-1 (Fig. 2f), indicative of a mechanistic link of vIRF-1 and NIX. This functional interaction was further assessed by transient transfection experiments; surprisingly, overexpression of both vIRF-1-Flag and V5-NIX, but not each individually, could induce a significant decrease in mitochondrial content, as determined by TFAM immunostaining (Fig. 2g). The decrease of mitochondria content was inhibited by autophagy/mitophagy inhibitors including leupeptin (a lysosomal protease inhibitor), liensinine (an inhibitor of autophagosome-lysosome fusion), and Mdivi-1 (a mitochondrial division inhibitor) (Supplementary Fig. 3), suggesting strongly that vIRF-1 and NIX together induce mitochondrial clearance by activating mitophagy. To rule out a role of nucleus-localized vIRF-1 in the mitochondrial clearance, we used the nuclear localization signal (NLS)-mutated version of vIRF-1 (NLS^X^) as previously described^24,25^. The vIRF-1 NLS^X^ variant strongly induced mitochondrial clearance together with V5-NIX (Fig. 2g). However, the vIRF-1 mutant (ΔPD) lacking the proline-rich domain (PD) that contains an atypical mitochondrial targeting signal sequence (MTS)^25^ could not induce mitochondrial clearance (Fig. 2g). These results suggest that mitochondrial localization of vIRF-1 is required for mitochondrial clearance. Since overexpression of NIX alone did not induce mitochondrial clearance (Fig. 2g), NIX may require an activation step to function as a mitophagy receptor. For example, it is known that in response to high oxidative phosphorylation activity, the small GTPase RHEB (Ras homolog enriched in brain) is recruited to the mitochondria and interacts with NIX to induce mitophagy^30^. Likewise, we hypothesized that mitochondria-localized vIRF-1 may activate NIX-mediated mitophagy by interacting with NIX.

**Fig. 3.**
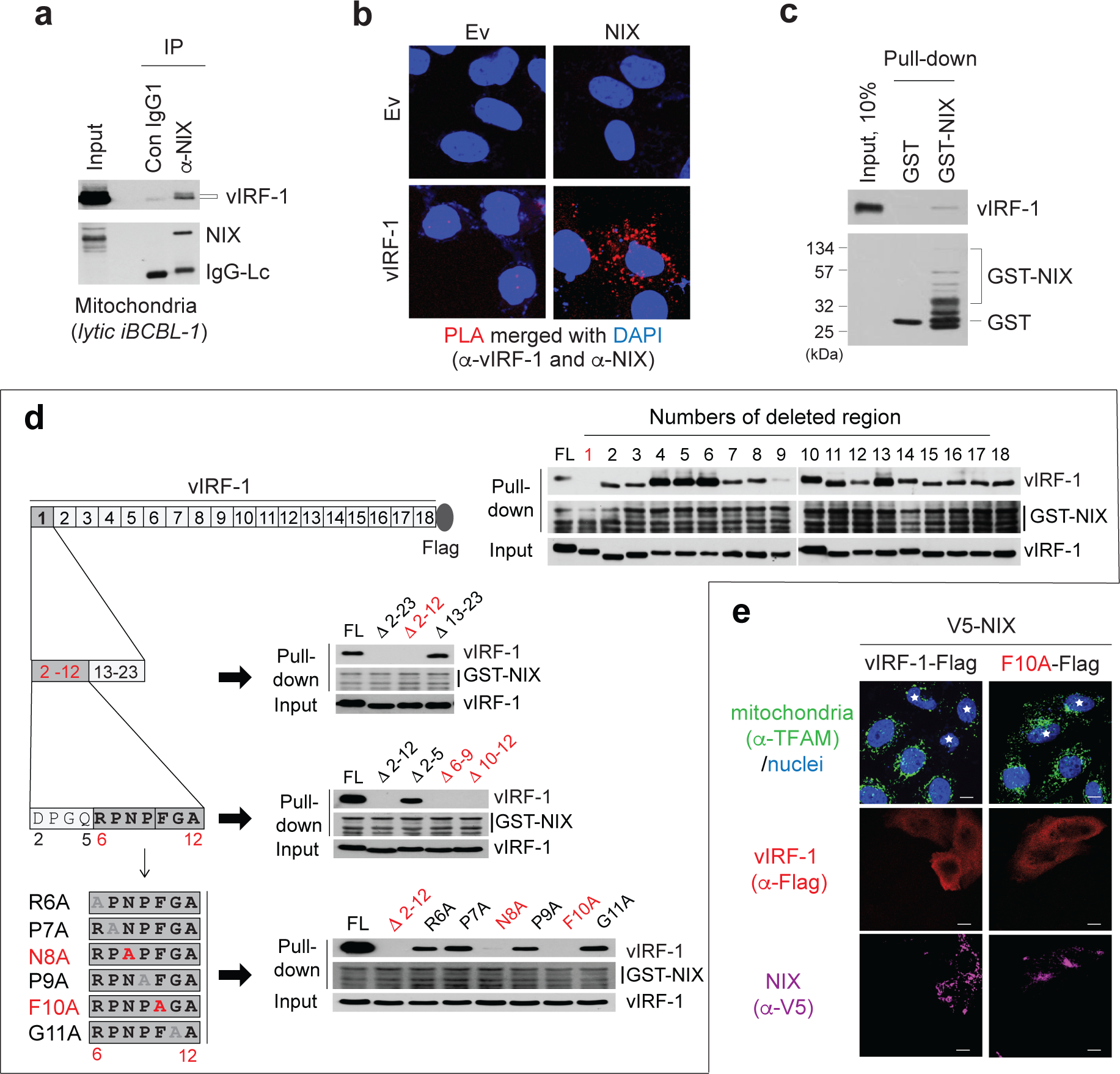
vIRF-1 interacts physically and functionally with NIX. **a** Detection of intracellular interaction of vIRF-1 and NIX in the mitochondrial fraction derived from iBCBL-1 cells, which were treated with Dox for 2 days, using co-immunoprecipitation (co-IP). Isotype-matched control antibody (Con IgG1) was used at an equivalent concentration. **b** *In situ* proximity ligation assay of vIRF-1 interaction with NIX in HeLa.Kyoto cells that were transfected with vIRF-1 and/or NIX. Rabbit anti-vIRF-1 and mouse anti-NIX (E-1) antibodies were used. See the SATR methods for further details. **c** *In vitro* binding assay of vIRF-1 interaction with NIX. Purified recombinant vIRF-1 was pulled down with glutathione sepharose beads coated with 1 µg of GST or GST-NIX. The pull-down complexes were separated by SDS-PAGE and immunoblotted with the indicated antibodies. **d** Identification of the vIRF-1 single residues required for NIX binding. Initial screening employed a series of vIRF-1 variants in which each of 18 segments of about 25 residues were deleted; next, the first N-terminal segment was progressively deleted. Finally, individual alanine substitutions of residues 6 to 11 were introduced. GST-pull down assays were performed with cell extracts of 293T cells expressing vIRF-1 full-length (FL) or variants. The regions or residues found to be required for NIX binding are highlighted in red. **e** IFA analysis of mitochondria content (TFAM) in HeLa.Kyoto cells that were transiently transfected for 24 h with V5-NIX and vIRF-1-Flag (wild type or F10A). The double transfected cells are marked with white stars. Scale bar, 5 µm.

### vIRF-1 binds directly to mitochondria-associated NIX

To examine whether vIRF-1 interacts with NIX on the mitochondria, we performed a co-immunoprecipitation assay with the mitochondrial fraction isolated from lytic iBCBL-1 cells. The result showed that vIRF-1 was indeed co-precipitated with NIX (Fig. 3a). Next, we employed a proximity ligation assay (PLA Duolink, Sigma) to analyze *in situ* interaction between vIRF-1 and NIX; their interaction was detected in the cytoplasm, potentially in mitochondria, of HeLa.Tokyo cells co-transfected with vIRF-1 and NIX, but not with each alone (Fig. 3b). To examine whether vIRF-1 binds directly to NIX, we performed an *in vitro* GST pull-down assay using purified recombinant proteins. The result showed that recombinant vIRF-1 protein, which was purified from bacteria as described previously^25^, was co-precipitated with purified GST-NIX but not GST protein (Fig. 3c).

To further characterize this interaction, we next sought to identify the vIRF-1 region(s) responsible for NIX binding. We used an *in vitro* binding assay using purified GST-NIX and cell lysates of 293T cells transfected with vIRF-1 full-length (FL) or a series of deletion mutants in which segments of 23 to 25 residues were deleted (Fig. 3d). Deletion of the first segment encompassing residues 1 to 23, comprising part of the PD region, completely abrogated NIX binding of vIRF-1 (Fig. 3d). Subsequent mapping and mutagenesis studies were carried out to identify vIRF-1 residues sufficient and required for NIX binding (Fig. 3d). These experiments identified two residues of vIRF-1, asparagine 8 (N8) and phenylalanine 10 (F10), as essential for NIX binding (Fig. 3d). Furthermore, the F10A mutation led to a loss of the ability of vIRF-1 to induce mitochondrial clearance in the presence of NIX (Fig. 3e), showing that vIRF-1 interaction with NIX is required to activate NIX-mediated mitophagy. Collectively, our results indicate that the N-terminal PD region of vIRF-1 contains motifs for vIRF-1 binding to and activating of NIX-mediated mitophagy and for mitochondrial targeting^25^.

### NIX determinants of physical and functional interactions with vIRF-1

To investigate the intracellular interaction of vIRF-1 and NIX, we employed a luciferase-based protein fragment complementation assay, NanoLuc^®^ Binary Technology (NanoBiT, Promega). The NanoBiT system consists of two small subunits, Large BiT (LgB; 17.6 kDa) and Small BiT (SmB; 1.3 kDa) (Fig. 4a). To find an optimal orientation of the NanoBiT tags for detection of vIRF-1:NIX interaction, Flag-vIRF-1 and V5-NIX were tagged at either the N- or C-terminal end of the NanoBiT subunits (Supplementary Fig. 4a). Intriguingly, when the NanoBiT tags were fused to the C-terminal end of NIX, the formation of a NIX dimer was greatly reduced (Supplementary Fig. 4b); this is likely due to its steric hindrance in the C-terminal tail anchor (TA) region, thus interfering with targeting to the OMM and/or TA region-mediated dimerization^31^. At 24 h after co-transfection with the different vIRF-1 and NIX plasmid combinations or single plasmids, a Nano luciferase (NLuc) substrate, furimazine, diluted in phosphate-buffered saline was added to the cell cultures. We identified two binary combinations showing the highest NLuc activity: 1) vIRF-1-SmB plus LgB-NIX and 2) SmB-vIRF-1 plus LgB-NIX (Supplementary Fig. 4c). To further demonstrate their specific interactions, we generated various deletion and substitution variants together with a negative control, HaloTag (HT)-SmB (Fig. 4b). NIX indeed bound specifically to vIRF-1; the luminescence signal was 10-fold higher when LgB-NIX was co-expressed with vIRF-1-SmB than HT-SmB (Fig. 4c). However, NIX binding to vIRF-1 was significantly diminished when the PD of vIRF-1 was deleted (Fig. 4d). These results are consistent with the data from the precipitation assays above and validated the NanoBiT system for further quantitative analysis of vIRF-1 and NIX interaction. We next examined whether the mitochondrial targeting of NIX is required for binding to vIRF-1. For this assay, we generated the NIX mutant (NIXΔTA) that is deficient in the tail anchor (TA) region (residues 188 to 208) and showed that this mutant lost the ability to localize to mitochondria and to form a homodimer (Supplementary Fig. 5 and Fig. 4e). The result showed that vIRF-1 indeed did not interact with NIXΔTA (Fig. 4f). To further confirm the requirement of NIX mitochondrial localization for vIRF-1 binding, we substituted the TA of NIX with that of other mitochondrial TA proteins, VAMP1B and FIS1. The results revealed that both the chimeric proteins, NIX-TA^VAMP1B^ and NIX-TA^FIS1^, properly localized to mitochondria (Supplementary Fig. 5) and could bind to vIRF-1 (Fig. 4f).

**Fig. 4.**
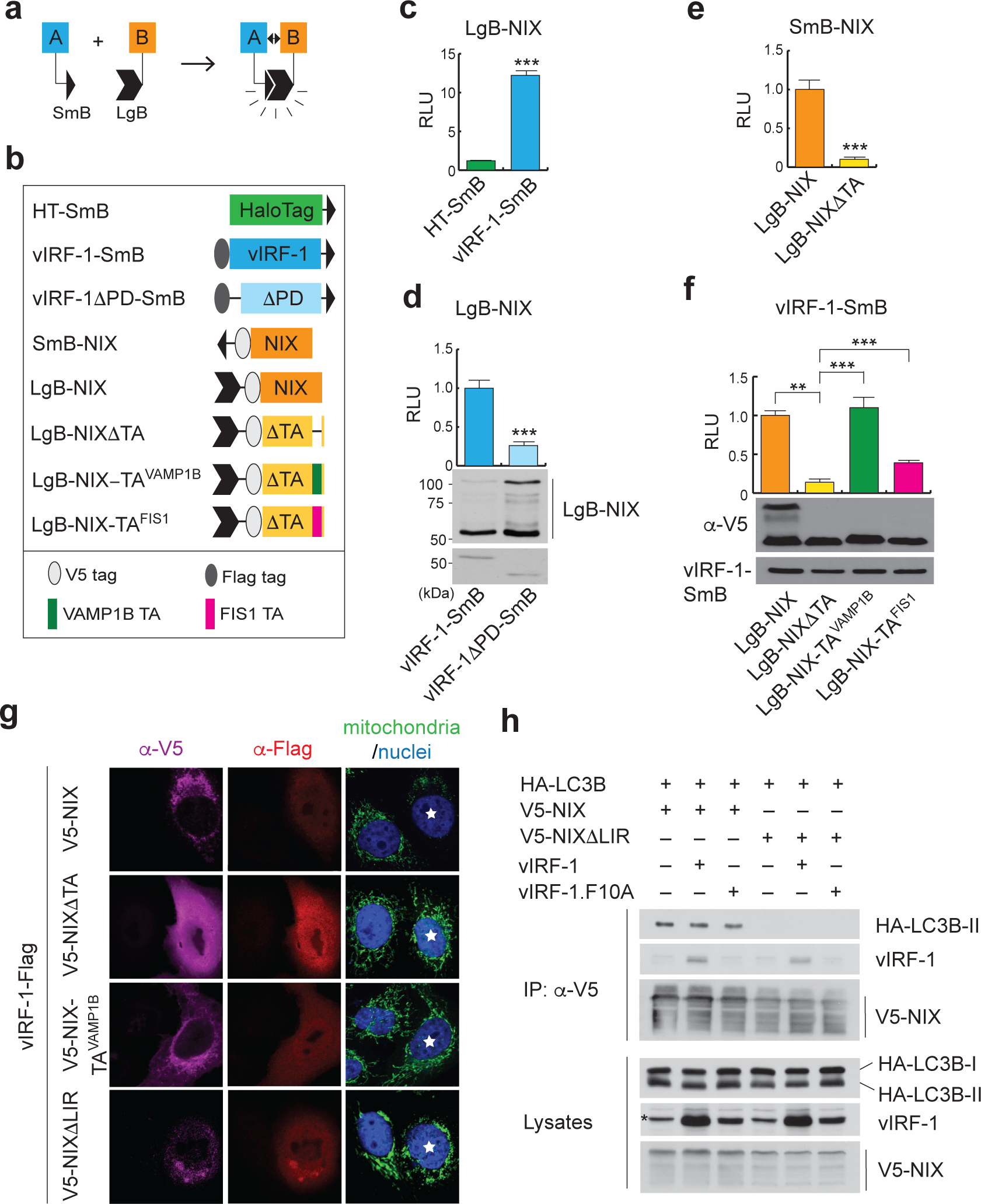
Characterization of NIX with respect to physical and functional interactions with vIRF-1. **a** Principle of the NanoBiT-based protein fragment complementation assay (PCA). When a protein [A] binds to its partner [B], their fused Small BiT (SmB; 1.3 kDa) and Large BiT (LgB; 17.6 kDa) fragments are brought into proximity, which allows structural complementation thus yielding a functional enzyme acting on Nano luciferase substrate. **b** Schematic structure of the NanoBiT-fused proteins used in the following studies. **c-f** NanoBiT PCA assays were performed using 293T cells transiently transfected with the indicated NanoBiT binary plasmids as described in the STAR methods. Each value represents the mean of triplicate samples from two independent experiments. Error bars represent standard deviation. (\*\**p* < 0.01 and \*\*\**p* < 0.001) **c** The optimal orientation of vIRF-1 and NIX (vIRF-1-SmB and LgB-NIX) was compared to HaloTag (HT)-SmB and LgB-NIX fusion proteins (See also Supplementary Fig. 4). **g** IFA analysis of mitochondria content (TFAM) in HeLa.Kyoto cells that were transiently transfected for 24 h with vectors expressing vIRF-1-Flag and V5-NIX (wild type or variants, NIX-TA^VAMP1B^ and NIXΔLIR lacking the LC3-interacting region (LIR)). Double-transfected cells are marked with white stars. **h** Co-IP analysis of the effect of vIRF-1 on NIX interaction with LC3B. HeLa.Kyoto cells transiently transfected with vectors expressing HA-LC3B and V5-NIX wild type or mutant (NIXΔLIR) in the presence or absence of vIRF-1 wild type and F10A mutant. Cells were treated overnight with bafilomycin A1 before their harvest. Asterisk indicates a non-specific band.

**Fig. 5.**
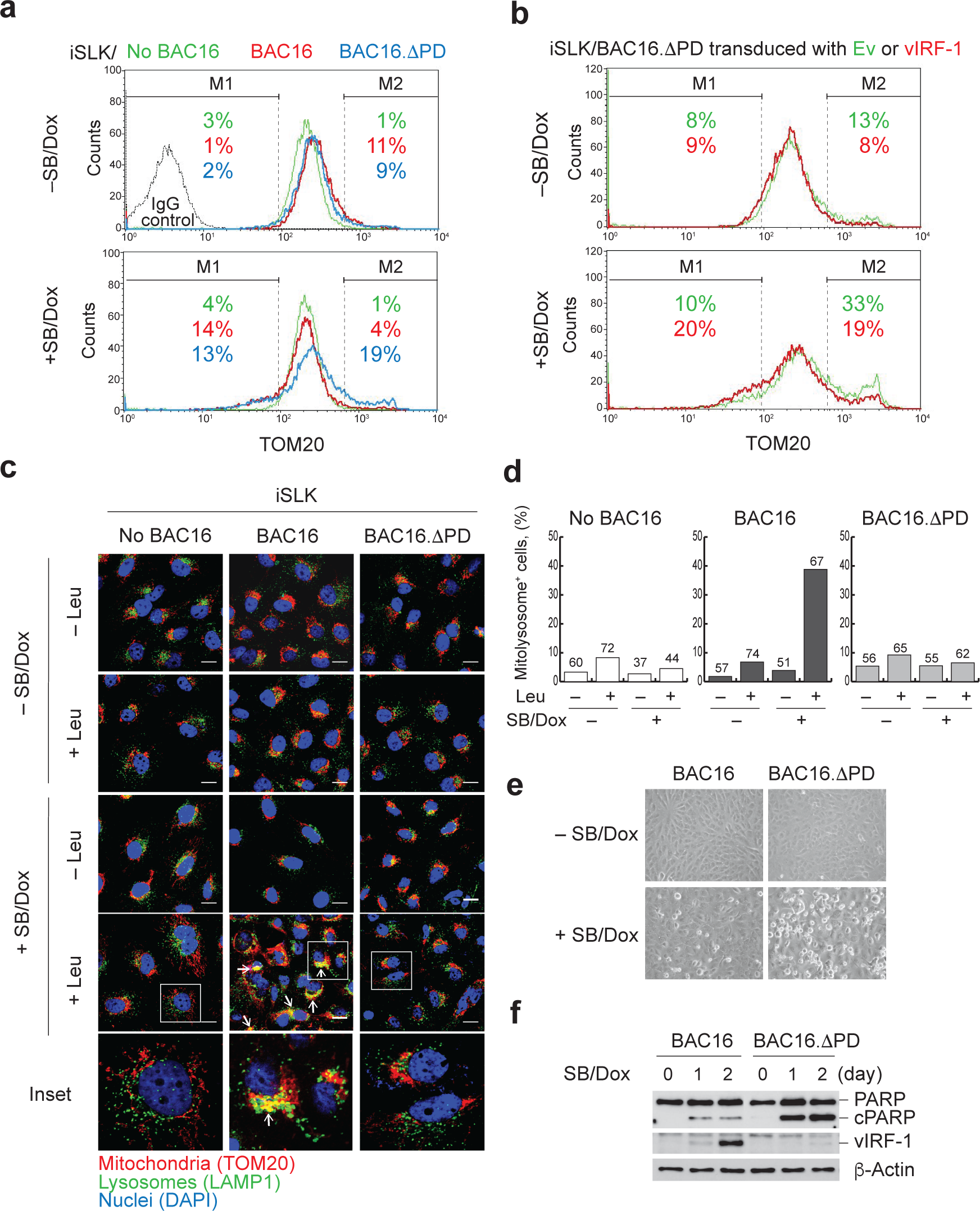
Mitochondrial targeting of vIRF-1 is essential for activating mitophagy and inhibiting cell death during virus replication. **a** Flow cytometric analyses of mitochondria content (TOM20) of iSLK cells that were uninfected or infected with HHV-8 BAC16 wild type or its variant BAC16.ΔPD that lacks the PD region of vIRF-1. See also Supplementary Fig. 6. The iSLK cells were left untreated or treated with sodium butyrate (SB) and Dox for 2 days. **b** iSLK BAC16.ΔPD cells were lentivirally transduced with control empty vector (Ev) or vIRF-1 and subjected to flow cytometric analysis as above. Histogram M1 and M2 regions indicate the populations with lower and higher levels of TOM20, respectively, compared to the peak area. **c, d** IFA analysis of mitolysosomes from iSLK, iSLK BAC16, and iSLK BAC16.ΔPD cells that were left untreated or treated with SB and Dox for 2 days. To facilitate detection of mitolysosomes, lysosomal protease inhibitor leupeptin (leu) or vehicle control (water) was added to the cultures 8 h before fixation. Scale bar, 10 µm. **d** Determination of the number of mitolysosome-containing cells. Multiple images were taken randomly of each sample from (c), and the number of cells showing apparent colocalization (i.e., yellow areas indicated by white arrows) of mitochondria (TOM20) and lysosomes (LAMP1) were counted and the derived data are presented graphically. **e** Phase-contrast images of iSLK BAC16 and BAC16.ΔPD cells untreated and treated with SB and Dox for 2 days. **f** Immunoblots of total cell extracts derived from iSLK BAC16 and BAC16.ΔPD cells treated with SB and Dox for 0, 1, and 2 days.

Unlike NIX, however, immunoblotting analysis showed that the chimeric NIX proteins did not form homodimers (Fig. 4f). Furthermore, NIX-TA^VAMP1B^ did not induce mitochondrial clearance together with vIRF-1 (Fig. 4g). These results indicate that NIX dimerization via its own TA region is essential for mitophagy activation, and the interaction of vIRF-1 and NIX is not sufficient for triggering mitophagy without NIX dimerization. Thus, we wondered how vIRF-1 activated NIX-mediated mitophagy. ATG8 binding to the LIR motif of NIX is critical in the NIX-mediated mitophagy pathway^17^. We thus generated the NIX mutant (NIXΔLIR) deficient in the LIR motif (residues 35 to 39) and found that vIRF-1 did not induce mitochondrial clearance together with NIXΔLIR (Fig. 4g). Accordingly, we postulated that vIRF-1 may promote the interaction of NIX with ATG8 family members via the LIR. To test this, we employed a co-IP assay using HA-tagged LC3B (HA-LC3B). However, we could not discern any effect of vIRF-1 on NIX and LC3B interaction while vIRF-1, but not vIRF-1 F10A, bound to NIX (Fig. 4h). These results suggest that vIRF-1 may activate NIX-mediated mitophagy by promoting NIX interaction with other member(s) of the ATG8 family or via unknown mechanism(s).

### Mitochondria-localized vIRF-1 promotes survival of lytic cells by activating mitophagy

Given that the PD of vIRF-1 contains the sequences for both mitochondrial targeting and NIX binding, we predicted that deletion of the PD would impede virus replication-induced mitophagy. To test this, we generated an HHV-8 bacterial artificial chromosome 16 mutant (BAC16.ΔPD) encoding vIRF-1 lacking the PD region, using λ-Red recombination techniques described previously^32^. We verified the deletion of the targeted region and the integrity of the mutated versus wild-type BAC16 DNA by sequencing and gel electrophoresis after AvrII or SpeI digestion (Supplementary Fig. 6). BAC16 (wild type) and BAC16.ΔPD DNAs were stably transfected into iSLK cells, which express Dox-inducible RTA and are broadly accepted as a model cell line for HHV-8 infection^33^. We first investigated mitochondrial content using flow cytometry analysis of TOM20. Lytic reactivation of the iSLK cells was induced by treatment with 1 mM sodium butyrate (SB) and 1µg/ml Dox. The results showed that the population with higher TOM20 (the M2 region of Fig. 5a) was increased by five-fold in lytic BAC16.ΔPD cells relative to lytic BAC16 cells while the population with lower TOM20 of both BAC16 and BAC16.ΔPD cells was increased upon lytic reactivation (the M1 region of Fig. 5a). Therefore, these data support our hypothesis that vIRF-1 plays a critical role in the downregulation of mitochondria content via its PD region. In line with this, lentiviral transduction of vIRF-1 into BAC16.ΔPD cells promoted an increase in the TOM20 high population (the M1 region) but a decrease in the TOM20 low population (M2 region) (Fig. 5b). Furthermore, we assessed the formation of mitolysosomes in latently and lytically-infected BAC16 and BAC16.ΔPD cells by immunostaining of TOM20 and LAMP1, a lysosome marker. We used the lysosomal protease inhibitor leupeptin (Leu) to facilitate the detection of mitolysosomes in the iSLK cells (Fig. 5c). The results showed that the formation of mitolysosomes was evident in lytic BAC16 cells, but not in control (BAC16-negative) and lytic BAC16.ΔPD cells (Figs. 5c and 5d). These genetic studies strongly support our hypothesis that mitochondria-targeted vIRF-1 activates mitophagy during HHV-8 lytic replication.

**Fig. 6.**
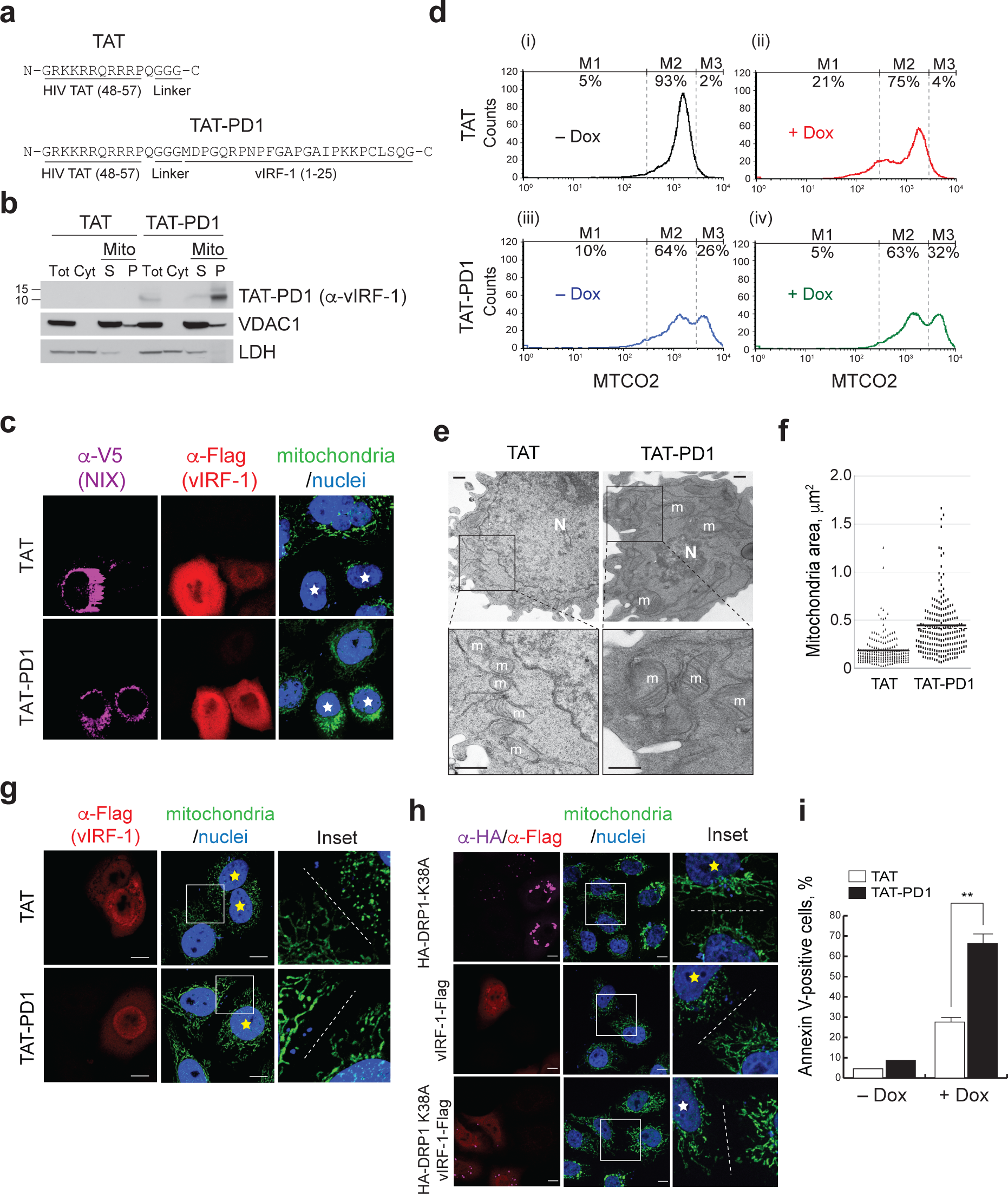
A cell-penetrating vIRF-1 peptide impairs mitophagy and mitochondrial dynamics. **a** Primary structures of TAT and TAT-PD1 peptides. PD1 consists of the N-terminal vIRF-1 residues 1 to 25. **b** Targeting of TAT-PD1 peptide to the mitochondrial detergent resistant (P, pellet) and, to a lesser extent, soluble (S) mitochondrial fractions of iBCBL-1 cells. VDAC1 and LDH were used for markers of mitochondria (Mito) and cytosol (Cyt), respectively. Tot, total cell extracts. **c** IFA analysis of mitochondrial content (TFAM) of HeLa.Kyoto cells that were transiently transfected with V5-NIX and vIRF-1-Flag and then treated with 20 µM of TAT or TAT-PD1 peptides. **d** Flow cytometric analysis of mitochondria content (MTCO2) of iBCBL-1 cells that were treated with 10 µM of TAT or TAT-PD1 peptides together with or without Dox for 2 days. **e, f** Electron microscopy analysis of lytic iBCBL-1 cells treated with 10 µM TAT or TAT-PD1 peptides together with Dox for 2 days. ‘m’ indicates mitochondria and ‘N’ indicates nucleus. Scale bar, 500 nm. Mitochondria areas were determined using ImageJ software and are depicted on the dot graph (f). The numbers of mitochondria and cells measured: n=187, 20 cells for TAT and n=208, 26 cells for TAT-PD1. **g** IFA analysis of mitochondria (TFAM) of HeLa.Kyoto cells transiently transfected with vIRF-1-Flag and treated with TAT or TAT-PD1 for 24 h. **h** IFA analysis of mitochondria (TFAM) of HeLa.Kyoto cells transiently transfected with a dominant negative form of DRP1 (HA-DRP1 K38A) and/or vIRF-1-Flag for 24 h. Images were taken at 63x magnification and a representative area containing mitochondria (inset) is expanded. Singly- and dually-transfected cells are highlighted in yellow and white stars, respectively. Dotted lines distinguish between transfected and untransfected cells. **i** Flow cytometric analysis of Annexin V binding to iBCBL-1 cells treated with 10 µM TAT or TAT-PD1 peptides together with or without Dox for 2 days. Each value represents the mean of two independent experiments and error bars represent standard deviation. (\*\**p* < 0.01).

We previously demonstrated that mitochondria-localized vIRF-1 plays an essential pro-survival role in lytic iBCBL-1 cells^25^. We thus wanted to confirm the results using BAC16-infected iSLK cells. When reactivated with SB/Dox treatment for 2 days, BAC16 iSLK cells exhibited modest morphological changes (cytopathic effects (CPE)) including rounding of cells, characteristic of apoptosis, as assessed by bright-field microscopy (Fig. 5e). However, BAC16.ΔPD cells showed stronger CPE (Fig. 5e). Furthermore, we observed that PARP cleavage, which is directly related to apoptotic cell death, was more apparent in BAC16.ΔPD cells than BAC16 cells upon reactivation (Fig. 5f). Taken together, these results suggest that mitochondria-localized vIRF-1 mediates, through promotion of mitophagic clearance of mitochondria, protection of lytic cells from mitochondrial-induced cytotoxicity.

### A cell-penetrating vIRF-1 PD peptide impairs mitophagy and mitochondrial dynamics

To test the possibility that a short peptide containing the mitochondrial- and NIX-targeting sequences of vIRF-1 may competitively inhibit vIRF-1-activated mitophagy, we synthesized a peptide (TAT-PD1) comprising vIRF-1 residues 1-25 fused to HIV-1 TAT residues 47-57 (Fig. 6a). The TAT-only peptide (hereafter termed TAT) was used as a control. Firstly, we examined whether TAT-PD1 can target to the mitochondria within cells. For immunoblotting detection of TAT-PD1, we used rabbit anti-vIRF-1 antibody, which recognizes an epitope in the N-terminus (residues 1 to 22). Consistent with our previous report^25^, a majority of TAT-PD1 could be detected in the mitochondrial fraction, in particular, the mitochondrial detergent resistant membrane fraction, but not the cytosolic fraction, of iBCBL-1 cells (Fig. 6b). Our IFA analysis showed that TAT-PD1, but not TAT, could inhibit the mitochondrial clearance induced by vIRF-1 and NIX in HeLa.Kyoto cells and led to an accumulation of the mitochondria at the perinuclear area in the cells (Fig. 6c). Next, we examined whether TAT-PD1 influences mitochondrial content in HHV-8-infected cells. Flow cytometric analysis of TOM20 showed that TAT-PD1, but not TAT, elicited an increase of the cell population with a higher mitochondrial content in lytic iBCBL-1 cells (compare M3 between Fig. 6d (ii) and (iv)). Also, TAT-PD1 effectively diminished the cell population with lower mitochondrial content (compare M1 between Fig. 6d (ii) and (iv)), indicating that TAT-PD1 could promote an accumulation of the mitochondria by blocking mitochondrial clearance in lytic cells. Intriguingly, TAT-PD1 also led to an accumulation of the M3 population in latent cells, but to a lesser extent compared to lytic cells (Fig. 6d (iii)), implying that TAT-PD1 might interfere with basal mitophagy, presumably mediated by vIRF-1 that is expressed at low levels in latent PEL cells or by a vIRF-1-independent mitophagy pathway.

Most strikingly, electron microscopy analysis revealed that TAT-PD, but not TAT alone, induced a substantial enlargement of the mitochondria in lytic iBCBL-1 cells (Figs. 6e and 6f). This may be attributed to an accumulation of fused mitochondria and/or inhibition of mitochondria fission by the peptide, together with concomitant inhibition of mitophagy. Interestingly, overexpression of vIRF-1 induced mitochondrial fragmentation, and TAT-PD1, but not TAT, inhibited vIRF-1-induced mitochondrial fragmentation (Fig. 6g). The fragmentation could be mediated by a block of fusion and/or promotion of fission. To test this, we used a dominant negative variant (K38A) of DRP1, a mitochondrial fission protein and found that vIRF-1-induced mitochondrial fragmentation was inhibited by the variant (Fig. 6h), suggesting that vIRF-1 may induce mitochondrial fragmentation potentially via DRP1. As mitochondria fission is often associated with initiation of mitophagy^34^, it is likely that vIRF-1-induced mitochondrial fission may contribute to the promotion of mitophagy although it remains to be determined the exact mechanisms by which vIRF-1 orchestrates mitochondrial dynamics to induce mitophagy. Interestingly, the electron microscopy image showed that TAT-PD1 induced nuclear condensation in lytic cells (Fig. 6e), indicative of apoptotic cell death. Supporting this observation, annexin V staining analysis showed that TAT-PD1 significantly promoted apoptotic cell death of lytic iBCBL-1 cells (Fig. 6i).

### Mitophagy is essential for HHV-8 productive replication

We previously demonstrated that mitochondria-localized vIRF-1 contributes to HHV-8 productive replication^3,25^. We sought to verify these findings using the Dox-inducible system described above. Consistent with our published data, production of encapsidated virions as well as expression of lytic antigens including ORF45 and K8.1 decreased by 40 to 50% in lytic vIRF-1-depleted iBCBL-1 cells compared to lytic control (sh-Luc) cells (Figs. 7a and 7b). Furthermore, the loss of the vIRF-1 function was exacerbated by simultaneous depletion of NIX (Figs. 7a and 7b). However, depletion of NIX alone had a marginal effect on the inhibition of virus productive replication and ORF45 expression. These results are reminiscent of our earlier mitochondrial content studies (Fig. 2f), suggesting that vIRF-1/NIX-mediated mitophagy is essential for successful lytic replication. Furthermore, we examined the effect of the mitophagy inhibitory peptide on virus productive replication. As expected, TAT-PD1, but not TAT, significantly inhibited the production of encapsidated HHV-8 virions in iBCBL-1 cells that were treated with Dox for 48 and 72 h (Fig. 7c). Interestingly, spontaneous virus production in the absence of Dox was greatly suppressed by TAT-PD1 (Fig. 7c), conceivably due to a cytotoxic effect caused by an accumulation of mitochondria as shown in Fig. 6d. Furthermore, a mitophagy inhibitor, liensinine, apparently inhibited basal and lytic production of HHV-8 virions at a final concentration of 20 µM (Fig. 7d), and a mitochondria fission inhibitor, Mdivi-1, also inhibited production of HHV-8 virions from lytic iBCBL-1 cells with a half maximal inhibitory concentration (IC_50_) of about 25 µM (Fig. 7e). Our findings demonstrate the importance of mitophagy, promoted by vIRF-1, for productive replication of HHV-8.

**Fig. 7.**
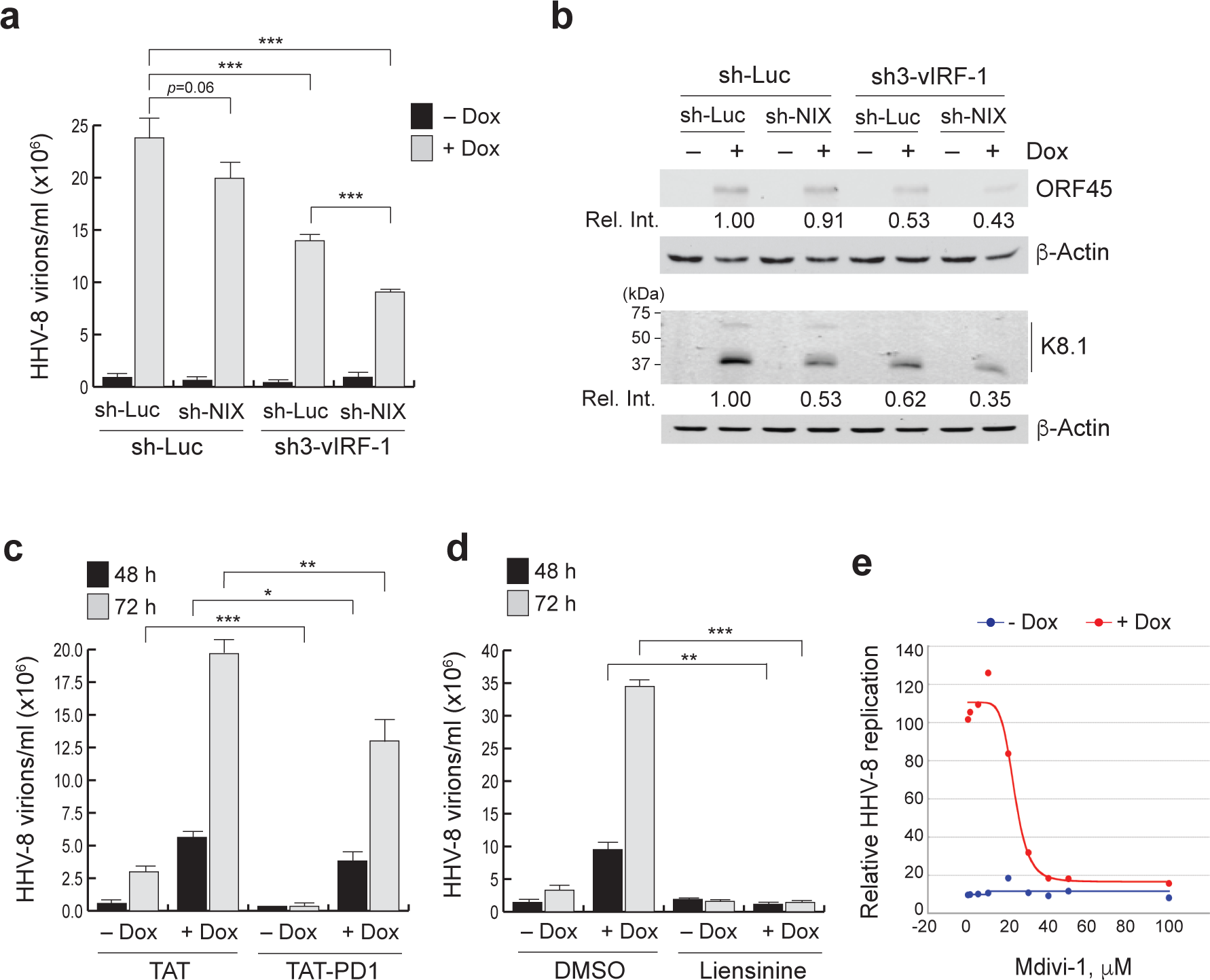
Mitophagy plays an essential role for HHV-8 productive replication. **a** qPCR analysis was employed to determine encapsidated viral genome copy numbers present in the media of control (sh-Luc) and vIRF-1-depleted iBCBL-1 cultures either untreated or treated with Dox for 4 days. Each value represents the mean ± standard deviation of a representative experiment performed in triplicate. (\*\*\**p* < 0.001) **b** Immunoblots of the total cell extracts derived from the same cultures as above (A) using antibodies to lytic antigens ORF45 and K8.1. The relative band intensities (Rel. Int.) of the antigens normalized to the loading control β-Actin are displayed beneath the corresponding panel. **c-e** qPCR analyses to measure encapsidated viral genome copy numbers in the media of iBCBL-1 cultures left untreated or treated with Dox for 2 days together with 10 µM TAT or TAT-PD1 peptides (c), 20 µM liensinine or vehicle (DMSO) (d), and Mdivi-1 (e) at the indicated concentrations. Each value represents the mean ± standard deviation from two or three independent experiments performed in triplicate. (\**p* < 0.05, \*\**p* < 0.01, and \*\*\**p* < 0.001)

## Discussion

The role of autophagy in HHV-8 lytic infection has been controversial. The HHV-8-encoded lytic proteins K7 and vBcl-2 suppress autophagy by interacting with autophagy proteins Rubicon and Beclin 1, respectively^35,36^, reportedly promoting virus replication. In addition, other lytic proteins including K1, vIL-6, vGPCR, and ORF45 may inhibit the initiation of autophagy by activating PI3K/AKT/mTOR signaling^37^. However, these studies are sharply in contrast to another report that found that RTA protein, a switch for turning on lytic reactivation, can activate autophagy to facilitate HHV-8 lytic replication^38^. A recent report also showed that autophagy is activated concurrently with HHV-8 lytic induction and is essential for viral lytic gene expression using multiple PEL cell lines^39^. Intriguingly, the authors proposed that the fusion of autophagosomes with lysosomes may be blocked at the late steps of virus replication to allow virus particles to be delivered to the extracellular milieu via autophagic vesicles. These results suggest that autophagy might be subject to different temporal regulation for HHV-8 productive replication. Nonetheless, the role of selective autophagy in HHV-8 infection has not previously been reported. Herein, we have provided experimental evidence for selective autophagy of mitochondria (mitophagy) in cells lytically infected with HHV-8, and further demonstrated that mitochondria-localized vIRF-1 activates mitophagy by directly interacting with the mitophagy receptor, NIX, to facilitate HHV-8 replication.

Our experiments with Dox-inducible double silencing of vIRF-1 and NIX in lytically HHV-8-infected PEL cells have demonstrated that vIRF-1 and NIX mechanistically linked for mitochondrial clearance. Conversely, overexpression of both vIRF-1 and NIX, but not each alone, promoted mitochondrial clearance in HeLa.Kyoto cells. This cooperation is likely to be mediated by vIRF-1 activation of NIX-mediated mitophagy rather than vice versa in that vIRF-1 utilized the LC3 interacting motif of NIX for mitochondrial clearance (Fig. 4g).

How does vIRF-1 activate NIX-mediated mitophagy? NIX forms a unique dimeric structure resistant to ionic detergents (e.g. sodium dodecyl sulfate) and reducing reagents (e.g. dithiothreitol) and its dimerization is functionally implicated in apoptosis and possibly mitophagy^40-42^. However, the precise details of the mechanisms by which dimerization contributes to the functions of NIX remain elusive. In fact, when overexpressed, NIX was able to readily dimerize but did not induce mitochondrial clearance by itself (Supplementary Fig. 2 and Fig. 2g). Moreover, co-transfected vIRF-1 induced mitochondrial clearance without promoting further dimerization of NIX. However, the NIX variants, NIX-TA^VAMP1B^ and NIX-TA^FIS1^, lost the ability to form a homodimer and to induce mitochondrial clearance when co-expressed with vIRF-1 although they could localize to mitochondria and bind to vIRF-1 (Fig. 4). Based on these data, NIX dimerization may be a prerequisite but not sufficient to mediate vIRF-1-activated mitophagy. A recent report showed that the phosphorylation of serine residues adjacent to the LIR of NIX stabilized the interaction with LC3B and promoted NIX-mediated mitophagy^43^. Because vIRF-1 did not enhance the interaction of NIX and LC3B, however, it is unlikely that vIRF-1 induces the NIX phosphorylation. Nevertheless, it is worth investigating in future studies if vIRF-1 promotes NIX interactions with the ATG8 family members other than LC3B. It is also conceivable that vIRF-1 activates other mitophagic pathways mediated by NIX. For example, NIX serves as a substrate of the E3 ubiquitin ligase PARK2, and then ubiquitinated NIX recruits the autophagy receptor NBR1 to the mitochondria^44^. Intriguingly, NIX was also shown to induce lysosomal degradation of mitochondria via MIEAP-induced accumulation of lysosome-like organelles within mitochondria, designated as MALM^45^. For future investigations, it would be interesting to examine if vIRF-1 also activates the non-canonical pathways of NIX for mitochondrial clearance.

In this study, we have also discovered a novel role of vIRF-1 in inducing mitochondrial fragmentation, potentially via a mitochondrial fission protein, DRP1, although the exact mechanism of action remains to be elucidated. Asymmetric mitochondrial fission, in which damaged or dysfunctional mitochondrial components are partitioned from intact components, has been postulated to play a key role in linking the altered mitochondrial components to the mitophagy machinery^46-48^. Interestingly, BNIP3 can induce both mitochondrial fragmentation and mitophagy^49,50^. Notably, DRP1 binding to BNIP3 and recruitment to mitochondria and mitochondrial fission appear to be a prerequisite for BNIP-mediated mitophagy in cardiomyocytes. Whereas NIX shares about 52% sequence identity (64% sequence similarity) with BNIP3, it is unclear whether NIX also plays a role in mitochondrial fission. Our data showed that overexpression of NIX often induced a clustering of mitochondria at the perinuclear area rather than fragmentation (Fig. 2g and Supplementary Fig. 5). Thus, it is likely that vIRF-1-induced mitochondrial fission may concomitantly contribute to the initiation of NIX-mediated mitophagy during virus replication.

It is currently unclear whether mitochondria are functionally altered and the damaged mitochondria trigger mitophagy during HHV-8 lytic replication. Here, we demonstrated that an accumulation of mitochondrial content could be cytotoxic to lytic cells and lead to amplification of antiviral responses including apoptosis and innate immune responses. Our findings suggest that mitochondria-localized vIRF-1 contributes to global regulation of the antiviral responses to HHV-8 lytic replication by activating NIX-mediated mitophagy, as well as specific regulation of the antiviral responses via its inhibitory interactions with proapoptotic BOPs and MAVS. Furthermore, this study identifies the vIRF-1-derived PD1 peptide as an anti-HHV-8 agent for treatment of HHV-8-associated diseases.

## Acknowledgement

We thank Dr. Edward Harhaj for critical reading of this manuscript. This work was supported by National Cancer Institute (NCI) grant R01CA214131 to Y.B.C. and by National Institute of Allergy and Infectious Diseases (NIAID) grant R21Al117168 to Y.B.C. This research was funded in part by a 2016-2017 developmental grant from the Johns Hopkins University Center for AIDS Research, an NIH funded program (P30AI094189 to Y.B.C.), and an NIH Shared Instrumentation grant (S10OD016374 to the JHU Microscope Facility).

## Author Contributions

M.T.V. designed the study, performed the experiments, and analyzed the results. B.J.S. performed the experiments. J.N. provided valuable suggestions, discussed the results, and proofread the manuscript. Y.B.C. designed the study, performed the experiments, analyzed the results, and wrote the manuscript.

## Materials & Correspondence

Young Bong Choi to whom correspondence and material requests should be addressed.

## Methods

### Cell culture

BCBL-1 (ATCC), TRExBCBL-1-RTA (a gift from Dr. Jae U. Jung, hereafter simply termed iBCBL-1), and iBCBL-1 derivative lines were cultured in RPMI 1640 medium (Quality Biological) supplemented with 15% fetal bovine serum (FBS), stable L-alanyl-glutamine (Glutamine XL), antibiotics including streptomycin and penicillin, and plasmocin prophylactic (Invivogen) at 37°C and 5% CO_2_. 293T, HeLa.Kyoto (a gift from Dr. Ron R. Kopito), and iSLK and iSLK-BAC16 (a gift from Dr. Jae U. Jung) cells were cultured in DMEM supplemented with 10% FBS, Glutamine XL, and antibiotics. Transient transfection with plasmids was performed using GenJet version II (SignaGen Laboratories) following the manufacturer’s instructions. For stable and doxycycline (Dox)-inducible expression of short hairpin RNAs (shRNAs), iBCBL-1 cells were lentivirally transduced with the indicated shRNAs in the presence of 10 µg/ml polybrene overnight and stably transduced cells were selected by growing in the presence of 1 µg/ml puromycin or 400 µg/ml geneticin for more than one month, and pooled clones were collected. Transfection of iSLK cells with bacmids BAC16 and BAC16.ΔPD was performed using Fugene^®^ HD (Promega) as described by the manufacturer’s protocol and the transfected cells were selected in the presence of hygromycin B (1,200 µg/ml) together with geneticin (250µg/ml) and puromycin (1 µg/ml). Reconstitution of iSLK/BAC16.ΔPD cells with vIRF-1 or empty vector was performed by lentiviral transduction in the presence of 10 µg/ml polybrene overnight.

### Lentivirus production

To produce lentiviruses, 293T cells were co-transfected with the lentiviral transfer vector together with the packaging plasmid psPAX2 and the vesicular stomatitis virus G protein expression plasmid pVSV-G at a ratio of 5:4:1. Two days later, virions were collected from the culture medium by ultracentrifugation in an SW28 rotor at 25,000 rpm for 2 h at 4°C. The virion pellets were resuspended in an appropriate volume of PBS to achieve 100x concentration. The transduction unit of lentiviruses was determined in 293T cells in the presence of appropriate antibiotics.

### Isolation of mitochondria

Pure mitochondria were isolated using Axis-Shield OptiPrep iodixanol (Sigma-Aldrich) as previously described ^51^. Briefly, latent and lytic iBCBL-1 cells were homogenized in buffer B (0.25 M sucrose, 1 mM EDTA, 20 mM HEPES-NaOH [pH 7.4]) with 50 strokes of a Dounce glass homogenizer and centrifuged at 1,000 × *g* for 10 min. An aliquot of homogenate was used as total cell extracts. The supernatant was further centrifuged at 13,000 × *g* for 10 min. The pellet was used as a crude fraction enriched in mitochondria. For further enrichment, the pellet was resuspended in 36% iodixanol, bottom-loaded under 10% and 30% iodixanol gradients, and centrifuged at 50,000 × *g* for 4 h. The mitochondria were collected at the 10%/30% iodixanol interface. For isolation of mitochondrial detergent-resistant membrane microdomains (mDRM), crude or pure mitochondria were incubated in TNE buffer (50 mM Tris-HCl [pH 7.4], 150 mM NaCl, and 1 mM EDTA) containing 1% Triton X-100 on ice for 30 min, and centrifuged at 21,000 × *g* for 10 min. The supernatant was used as a soluble mitochondrial fraction, and the pellet was used as the mDRM fraction. The pellet was boiled in 1× SDS sample buffer.

### DNA manipulation

All polymerase chain reaction (PCR) amplification and site-directed mutagenesis including point and deletion mutations were performed using Platinum™ *Pfx* or SuperFi™ DNA polymerase (Thermo Fisher Scientific). Subcloning of open reading frames (ORFs) and their derivatives into expression plasmids including pICE (a gift from Steve Jackson; Addgene plasmid #46960), pGEX-4T-1 (GE Healthcare Life Sciences), and NanoBiT^®^ system vectors (Promega) was performed using appropriate restriction enzyme sites (Supplementary Table 1). For replacement of the NIX tail-anchor (TA) region with the TA region of other tail-anchored proteins, overlap extension PCR was carried out as previously described^52^ and the fused genes were cloned into vectors pBiT1.1-N (Promega) and pICE_V5^32^.

### Mutagenesis of HHV-8 bacmid BAC16

The BAC16 genome was edited using a two-step seamless Red recombination in the context of *E. coli* strain GS1783 (a kind gift from Greg Smith) as previously described^53^. Briefly, PCR amplification was performed to generate a linear DNA fragment containing a kanamycin resistance expression cassette, an I-SceI restriction enzyme site, and flanking sequences derived from HHV-8 genomic DNA, each of which includes a 40-bp copy of a duplication. The direct connection of the translational initiation codon with codon 76, which results in deletion of the proline-rich domain (PD, 2 to 75 residues) of vIRF-1, was placed in the middle of the duplication (Supplementary Fig. 6a). This fragment was purified and then electroporated into GS1783 cells harboring BAC16 and transiently expressing *gam, bet*, and *exo*, which are expressed in a temperature-inducible manner from the lambda Red operon in GS1783 chromosome. The integrated Kan^R^/I-SceI cassette was cleaved by I-SceI enzyme that was inducibly expressed by treatment with 1% L-arabinose, resulting in a transiently linearized BAC16. A second Red-mediated recombination between the duplicated sequences results in recirculation of the BAC DNA and seamless loss of the Kan^R^/I-SceI cassette. Kanamycin-sensitive colonies were selected via replica plating. The BAC DNAs were purified using the NucleoBond BAC 100 kit (Clontech). The recombination area was amplified by PCR and the mutation was verified by DNA sequencing of the PCR amplicon (Supplementary Fig. 6a). Gross genome integrity and the deletion of the PD region were verified using digestion with AvrII and SpeI restriction enzymes and agarose gel analysis of digestion profiles (Supplementary Fig. 6b).

### Immunological assays

Antibodies used in immunological assays including immunoblotting, immunoprecipitation, and immunostaining are listed in Supplementary Table 2. For the preparation of total cell extracts, cells were resuspended in RIPA buffer (50 mM Tris [pH 7.4], 150 mM NaCl, 1% Igepal CA-630, and 0.25% deoxycholate) containing a protease inhibitor cocktail and protein phosphatase inhibitors including 10 mM NaF and 5 mM Na_3_VO_4_ and sonicated using Bioruptor (Diagenode) for 5 min in ice water at a high-power setting (320 W). For immunoblotting, cell lysates were separated by SDS-PAGE, transferred to nitrocellulose or polyvinylidene difluoride membranes, and immunoblotted with appropriate primary antibodies diluted in SuperBlock™ (phosphate-buffered saline [PBS]) blocking buffer (Thermo Fisher Scientific). Following incubation with horse radish peroxidase-labeled appropriate secondary antibody, immunoreactive bands were visualized by an enhanced chemiluminescent (ECL) reagent on an ECL film. ImageJ software (NIH) was used to quantify the signal from immunoblots. For immunoprecipitation (IP), total cell or mitochondrial extracts were incubated with specific primary antibody at 4°C overnight and incubated with protein G-agarose beads (Cell Signaling Technology) for an additional 3 h. Immunoprecipitants were washed with RIPA buffer, followed by elution of bound proteins with 1× SDS sample buffer. To avoid detection of IgG used in IP assays, we used Clean-Blot IP detection reagent (Thermo Fisher Scientific) was used. For indirect immunofluorescent assay (IFA), cells grown on a coverslip (and transfected) were fixed in Image-iT™ fixative solution (Thermo Fisher Scientific) and permeabilized in 0.5% Triton X-100 prepared in PBS. iBCBL-1 cells were attached on poly-L-lysine coated coverslips. Following incubation with SuperBlock™ PBS blocking buffer for 1 h at room temperature, coverslips were incubated with primary antibodies, washed with PBS, and then incubated with appropriate fluorescent dye-conjugated secondary antibodies. Coverslips were mounted in ProLong™ Gold Antifade Mountant containing DAPI (Thermo Fisher Scientific) on glass slides and cells were imaged on a Zeiss 700 confocal laser scanning microscope (LSM) with a 40x or 63x oil-immersion objective and Zen software. ImageJ was used for all image preparation and analysis.

### Quantification of mitochondrial DNA

The amount of mitochondrial DNA (mtDNA) in lytic iBCBL-1 cells was determined using a combined approach of IFA and fluorescence in situ hybridization (FISH). Firstly, iBCBL-1 cells were treated with Dox for 2 days and IFA was performed with rabbit anti-vIRF-1 antibody as described above to identify lytically infected iBCBL-1 cells. After incubation with an anti-rabbit IgG Alexa 594-conjugated secondary antibody, the cells were post-fixed in Image-iT™ fixative solution for 15 min in the dark. After washing in PBS three times, the cells were incubated with 0.1 µg/µl RNase A for 1 h at 37°C, permeabilized in ice-cold 0.7% Triton X-100 and 0.1M HCl for 10 min on ice, and denatured in 50% formamide and 2x SSC at 80°C for at least 30 min. Coverslips were dehydrated by successive rinsing in 70%, 80%, and 95% ethanol and air-dried, and then hybridized with mtDNA-specific probes overnight at 37°C in a dark and humid chamber. The probes were prepared by nick translation of six overlapping PCR products (see (Supplementary Table 3 for primer sequences) representing the entire human mtDNA using FISH Tag™ DNA green kit (Thermo Fisher Scientific). Following washing, coverslips were mounted, and cells were imaged as above. The amount of mtDNA were measured using ImageJ.

### Image and flow cytometric analyses of mitochondrial content

For cytometric analyses of mitochondrial content, cells were immunostained in suspension with antibodies to mitochondrial proteins including MTCO2 and TOM20. Briefly, cells were fixed in 4% formaldehyde at 37°C for 10 min and permeabilized in a final concentration of 90% methanol. The fixed cells were washed twice with blocking buffer (0.5% BSA in PBS) by centrifugation at 3,000 x g for 5 min and incubated with the indicated primary antibody in blocking buffer for 1 h at room temperature. Cells were washed twice with blocking buffer and stained with Alexa Flour^®^ 488-conjugated goat anti-mouse IgG antibody in blocking buffer. Image and flow cytometric analyses of the immunostained cells were performed using Cellometer Vision CBA Image Cytometer (Nexcelom) and FACSCalibur (BD Biosciences), respectively.

### Assessment of formation of mitochondria-containing lysosomes (mitolysosomes)

For assessment of mitolysosome formation in iBCBL-1 cells, mitochondria and lysosomes were labeled using CellLight^®^ reagent BacMam 2.0 (Thermo Fisher Scientific) as described in the manufacturer’s protocol. Briefly, iBCBL-1 cells were left untreated or treated with Dox and 24 h later infected with baculoviruses encoding CellLight^®^ Lysosomes-GFP and CellLight^®^ Mitochondria-RFP at a ratio of 30 particles per cell. Twenty-four hour later, cells were fixed and visualized on the Zeiss 700 confocal LSM. For assessment of mitolysosome formation in BAC16 iSLK cells, IFA was performed using antibodies to TOM20 and LAMP1 along with respective secondary antibodies conjugated to Alexa Flour^®^ 594 and Alexa Fluor^®^ 647 as described above.

### Electron microscopy

Latent and lytic iBCBL-1 cells were fixed in buffer containing 2.5% glutaraldehyde, 3 mM MgCl_2_, 0.1 M sodium cacodylate [pH 7.2] for 1 h at room temperature. After rinsing, samples were post-fixed in buffer containing 1% osmium tetroxide, 0.8% potassium ferrocyanide, 0.1 M sodium cacodylate for 1 h on ice in the dark. After rinsing with 0.1 M maleate buffer [pH6.2] at 4°C, samples were stained in 2% uranyl acetate in 0.1 M maleate buffer at 4°C for 1 h in the dark, dehydrated in a graded series of ethanol and embedded in Eponate 12 (Ted Pella) resin, and polymerized at 60°C overnight. Thin sections, 60 to 90 nm, were cut with a diamond knife on the Reichert-Jung Ultracut E Ultramicrotome and picked up with 2×1 mm formvar coated copper slot grids. Grids were imaged on a Philips CM120 at 80 kV and captured with an AMT XR80 CCD (8 megapixel) camera.

### Proximity ligation assay

Proximity ligation assay (PLA) was performed using the DuoLink^®^ *In Situ* Red Starter Kit Mouse/Rabbit (Sigma-Aldrich) following the manufacturer’s instruction. Briefly, HeLa.Kyoto cells were seeded at low density on glass coverslips and 24 h later transfected with vIRF-1 and/or NIX plasmids. The next day, the cells were fixed in Image-iT™ fixative solution and permeabilized in 0.5% Triton X-100 prepared in PBS for 15 min. Samples were incubated with 3% bovine serum albumin (BSA) for 1 h at 37°C in a humidified chamber and then overnight at 4°C with rabbit anti-vIRF-1 and mouse anti-NIX (H-8, Santa Cruz Biotechnology) antibodies. Slides were then incubated for 1 h at 37°C with a mix of the MINUS (anti-mouse) and PLUS (anti-rabbit) PLA probes. Hybridized probes were ligated using the Ligation-Ligase solution for 30 min at 37°C and then amplified using Amplification-Polymerase solution for 100 min at 37°C. Slides were then mounted using DuoLink^®^ II Mounting Medium with DAPI and imaged on the Zeiss 700 confocal LSM.

### GST-pull down assays

Bacterially expressed recombinant glutathione-S-transferase (GST) and GST-fusion NIX (GST-NIX) proteins were purified by standard methods. 1 µg GST or GST-NIX was incubated with 20 µl bed volume of washed glutathione sepharose-4B beads for 1 h at room temperature. After washing in binding buffer (PBS plus 1% Triton X-100), the protein-bead complexes were incubated with 1 µg purified recombinant vIRF-1 protein as described previously^25^, or 293T cell extracts containing vIRF-1-Flag proteins at 4°C overnight, washed in binding buffer four times, separated on SDS-PAGE, and subjected to immunoblotting.

### Protein fragment complementation assay (PCA)

PCA was performed using NanoBiT^®^ Protein:Protein Interaction System (Promega). Twenty-four hours after transfection of 293T cells in a 6-well plate with the indicated genes in the NanoBiT vectors, cells were washed in PBS and resuspended in 1 ml of Opti-MEM™ I reduced serum medium (Thermo Fisher Scientific), and 100 µl of the cell suspension was transferred to a 96-well white plate in triplicate. Furimazine (N1110, Promega), a NanoLuc^®^-luciferase substrate, was diluted in PBS at a ratio of 1 to 50 and 25 µl of the diluted substrate was added to each well. After 5 min of incubation, the luciferase activity in each well was measured by GloMax^®^ 96 microplate luminometer (Promega).

### Apoptosis assays

Dead cells existing prior to the experiments were removed using Histopaque-1077 (Sigma) or Dead Cell Removal kit (Miltenyl Biotec). Apoptotic cells were identified by staining with FITC-annexin V as described by the manufacturer’s protocol (BioLegend). Also, apoptotic cell death was monitored by detecting the cleaved product of PARP1 via immunoblotting.

### Real time-quantitative PCR (RT-qPCR) analysis of gene expression

Total RNAs were isolated using the RNeasy mini kit (Qiagen). First-strand cDNA was synthesized from 1 µg of total RNA using SuperScript IV reverse transcriptase (Thermo Fisher Scientific) with random hexamers. RT-qPCR was performed using an ABI Prism 7500 system (Applied Biosystems) with the FastStart SYBR green/ROX master mix (Sigma-Aldrich). Reactions were performed in a total volume of 25 µl and contained 50 ng of reverse-transcribed RNA (based on the initial RNA concentration) and gene-specific primers. PCR conditions included an initial incubation step of 2 min at 50°C and an enzyme heat activation step of 10 min at 95°C, followed by 40 cycles of 15 seconds at 95°C for denaturing and 1 min at 60°C for annealing and extension.

### HHV-8 replication assays

For determination of encapsidated HHV-8 genome copy number, viral DNA was purified using Quick-DNA™ Viral Kit (Zymo Research) following pretreatment of the culture supernatants containing HHV-8 virions with DNase I (New England BioLabs) at 37°C overnight. RT-qPCR was performed as described above with LANA primers (Supplementary Table 3). BAC16 DNA was used as a standard for the calibration curve. Also, the expression levels of lytic antigens including ORF45 and K8.1 were determined by immunoblotting.

### Reagents

Chemical reagents were purchased from the following companies: MG132, Cell Signaling Technology; doxycycline, bafilomycin A1, chloroquine, and sodium butyrate, MilliporeSigma; Liensinine, AK Scientific; Mdivi-1, Cayman Chemical. TAT and TAT-PD1 peptides were custom synthesized by Biomatik, Canada.

### Quantification and Statistical Analysis

Statistical parameters including statistical analysis, statistical significance, and *p* value are stated in the Figure legends and Supplemental Figure legends. Statistical analyses were performed using Synergy software (KaleidaGraph). Differences between controls and samples were determined by matched pair t-test and were considered significant when the *p* value was less than 0.05 (*p* < 0.05).

