## Supplementary Info for "Activation of NIX-mediated mitophagy and replication by an interferon regulatory factor homologue of human herpesvirus"

**Supplementary Information**

**Supplementary Fig. 1. Selective autophagy of mitochondria is activated in lytically HHV-8-infected cells.**

**a** IFA analysis of TOM20 and vIRF-1 in lytic iBCBL-1 cells that were treated with Dox for 2 days. vIRF-1-expressing cells are outlined by white dotted lines. Scale bar, 10 µm. **b** MitoTracker® Red staining of the mitochondria of lytic iBCBL-1 cells that were treated with Dox for 2 days. MitoTracker® Red was added to the culture 30 min before fixation, and then IFA was performed using anti-vIRF-1 antibody. vIRF-1-expressing cells are outlined by white dotted lines. **c, d** Image cytometric analysis of MTCO2 expression in lytic iBCBL-1 cells that were treated with Dox for 2 days. Autophagy inhibitors, bafilomycin A1 (Baf A1, 200 nM) (c) or leupeptin (40 µM) (d), or vehicle control (DMSO), was added to the cultures 6 h before fixation. **e** Immunoblots of total-cell extracts derived from iBCBL-1 cultures left untreated or treated with Dox for 2 days. The indicated inhibitors were added to the culture 6 h before harvesting the cells: 10 µM MG132, 50 µM chloroquine (CQ), and 200 nM Baf A1. The relative band intensities (Rel. Int.) of MTCO2 normalized to the loading control lactate dehydrogenase (LDH) are displayed beneath the corresponding panel.

**Supplementary Fig. 2. vIRF-1 has no effect on NIX dimerization.**

Immunoblot analysis of total-cell and mitochondrial extracts derived from 293T cells transiently transfected with the indicated amounts of V5-NIX plasmid in the presence or absence of vIRF-1-Flag. HSP60 was used as loading control. $(V5\text{-}NIX)_2$ indicates a homodimer form of NIX.

**Supplementary Fig. 3. Mitochondrial clearance induced by vIRF-1 and NIX is inhibited by autophagy/mitophagy inhibitors.**

IFA analysis of mitochondria content (TFAM) in HeLa.Kyoto cells that were transiently transfected with vIRF-1-Flag and V5-NIX. The following inhibitors and vehicle control (DMSO) were incubated with the transfected cells for 18 h: leupeptin (40 μM), liensinine (20 μM) and Mdivi-1 (20 μM). White asterisks indicate cells expressing both vIRF-1-Flag and V5-NIX.

**Supplementary Fig. 4. A combinatorial approach to determine the optimal orientation of NanoBiT-fused vIRF-1 and NIX proteins.**

**a** Diagram of NanoBiT (LgB or SmB)-fused Flag-vIRF-1 and V5-NIX at the N- and C-terminal ends. **b** Immunoblot analyses of extracts of 293T cells transiently transfected with the NanoBiT vectors as shown in (a) for expression analysis. Red asterisk indicates a homodimer form of NIX. **c** Measurement of luminescence intensity from 293T cells transfected with a single plasmid encoding NanoBiT-fused vIRF-1 or NIX or with eight combinations of the plasmids. Each value represents the mean ± standard deviation from a representative experiment. The binary sets providing the highest luminescence intensity are highlighted in red.

**Supplementary Fig. 5. Chimeric NIX-TA fused to heterologous tail-anchor domains can localize to mitochondria.**

IFA analysis of mitochondria localization of V5-NIX wild type and its variants including the deletion mutant (ΔTA) lacking the tail-anchor (TA) domain and chimeric constructs in which the native TA was replaced with the TA domains of other mitochondrial proteins including VAMP1B and FIS1. HeLa.Kyoto cells were transiently transfected with the indicated plasmids for 24 h. TOM20 was used as a marker of mitochondria. Scale bar, 10 μm.

**Supplementary Fig. 6. Generation and verification of the BAC16 mutant (BAC16.ΔPD) lacking the PD region of vIRF-1.**

**a** The targeted deletion of the coding sequences for vIRF-1 PD (residues 2 to 75) in the BAC16 DNA was performed using λ-Red-mediated recombination (see the STAR methods for details) and confirmed by DNA sequencing. **b** The genomic integrity of BAC16.ΔPD was verified by restriction analysis using AvrII and SpeI enzymes and comparison of the product profiles with those of wild-type BAC16. The diagrams show the cleavage sites of the enzymes on the BAC16 and BAC16.ΔPD bacmid DNAs. The digested DNAs were run on a 0.4% agarose gel at 0.6 V/cm and visualized by ethidium bromide staining (right panel). Blue and red arrows indicate the DNA fragments harboring the coding regions for vIRF-1 wild type and ΔPD mutant, respectively.

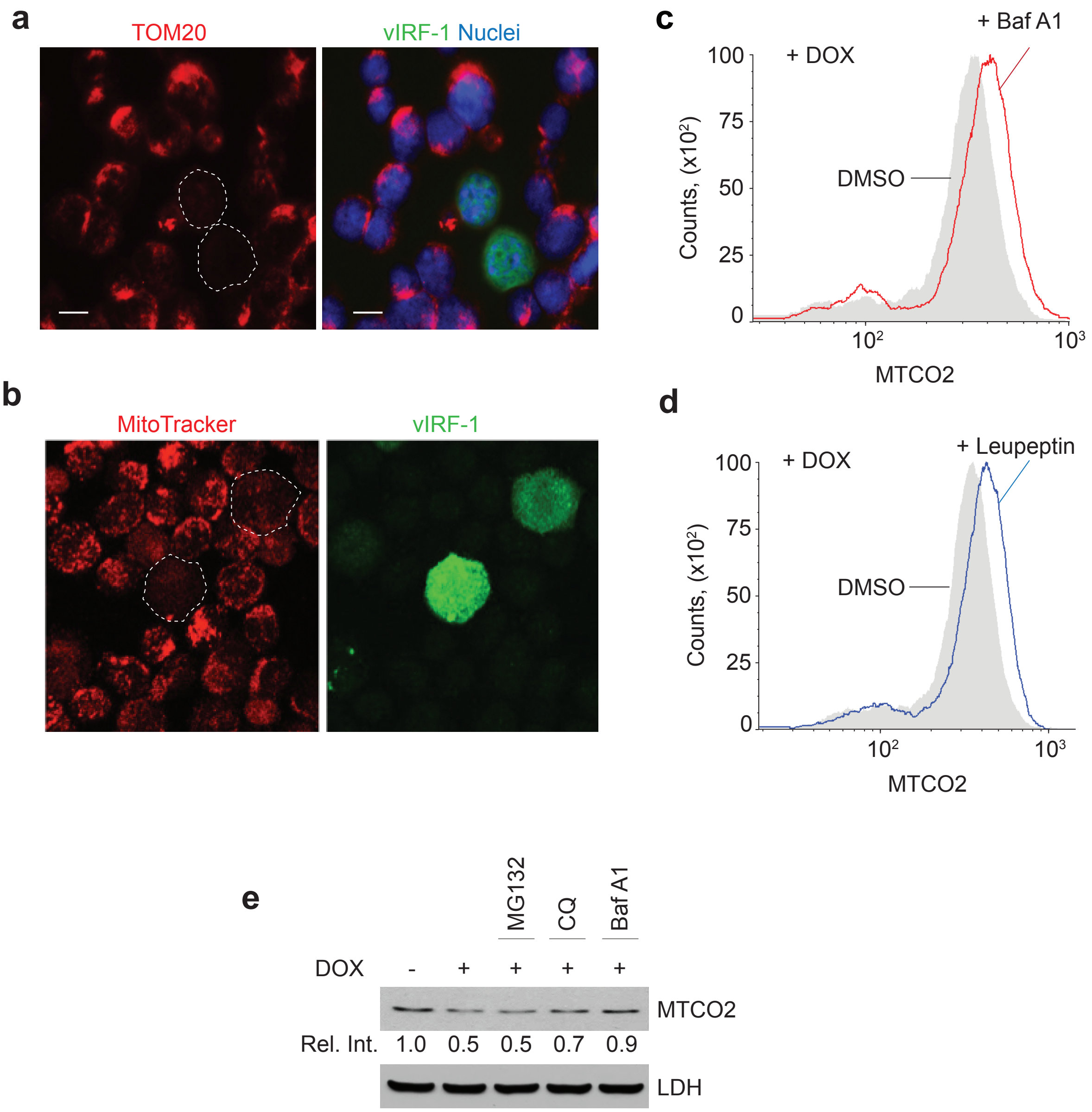

Supplementary Fig. 1

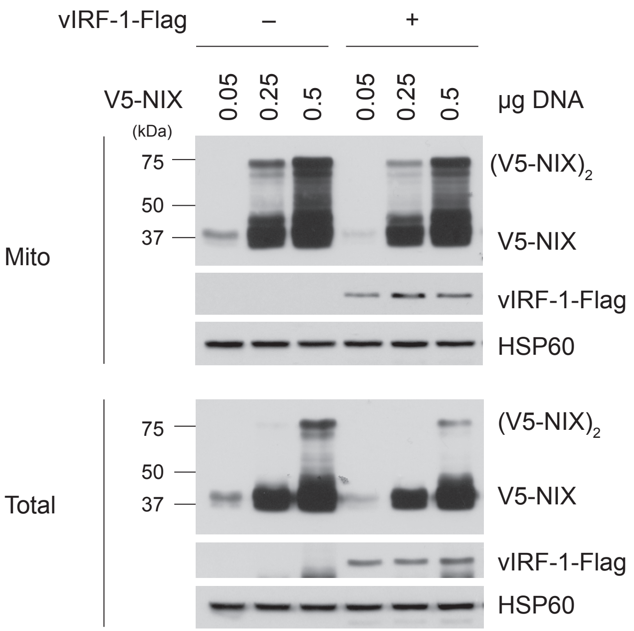

Supplementary Fig. 2

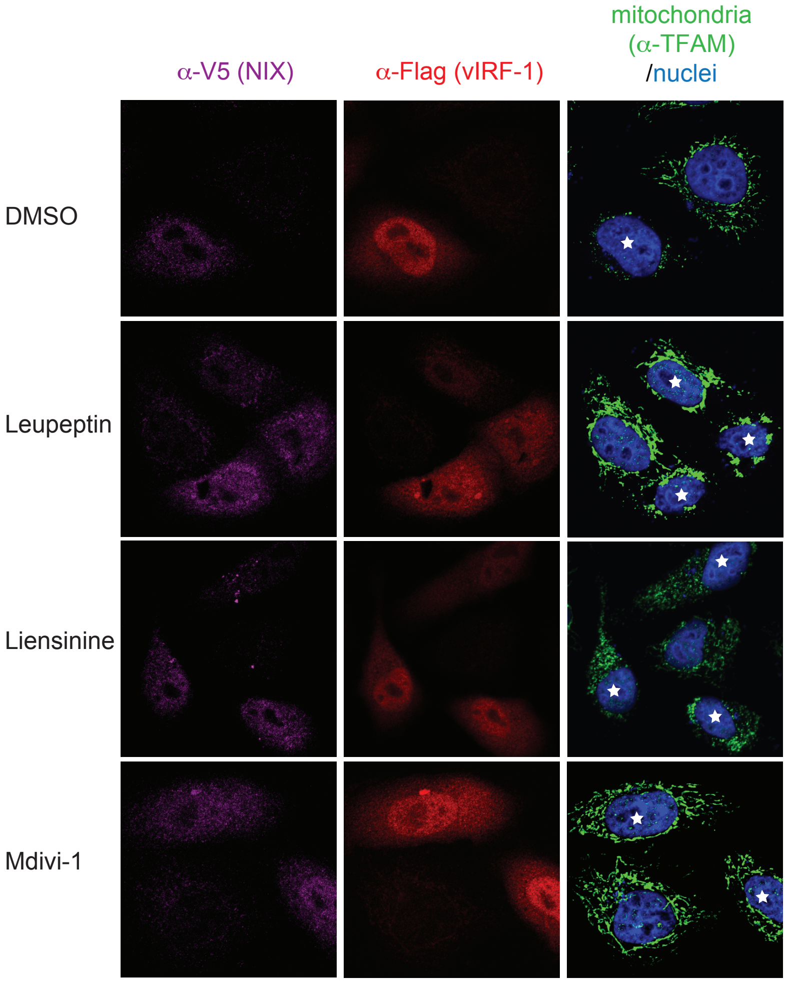

Supplementary Fig. 3

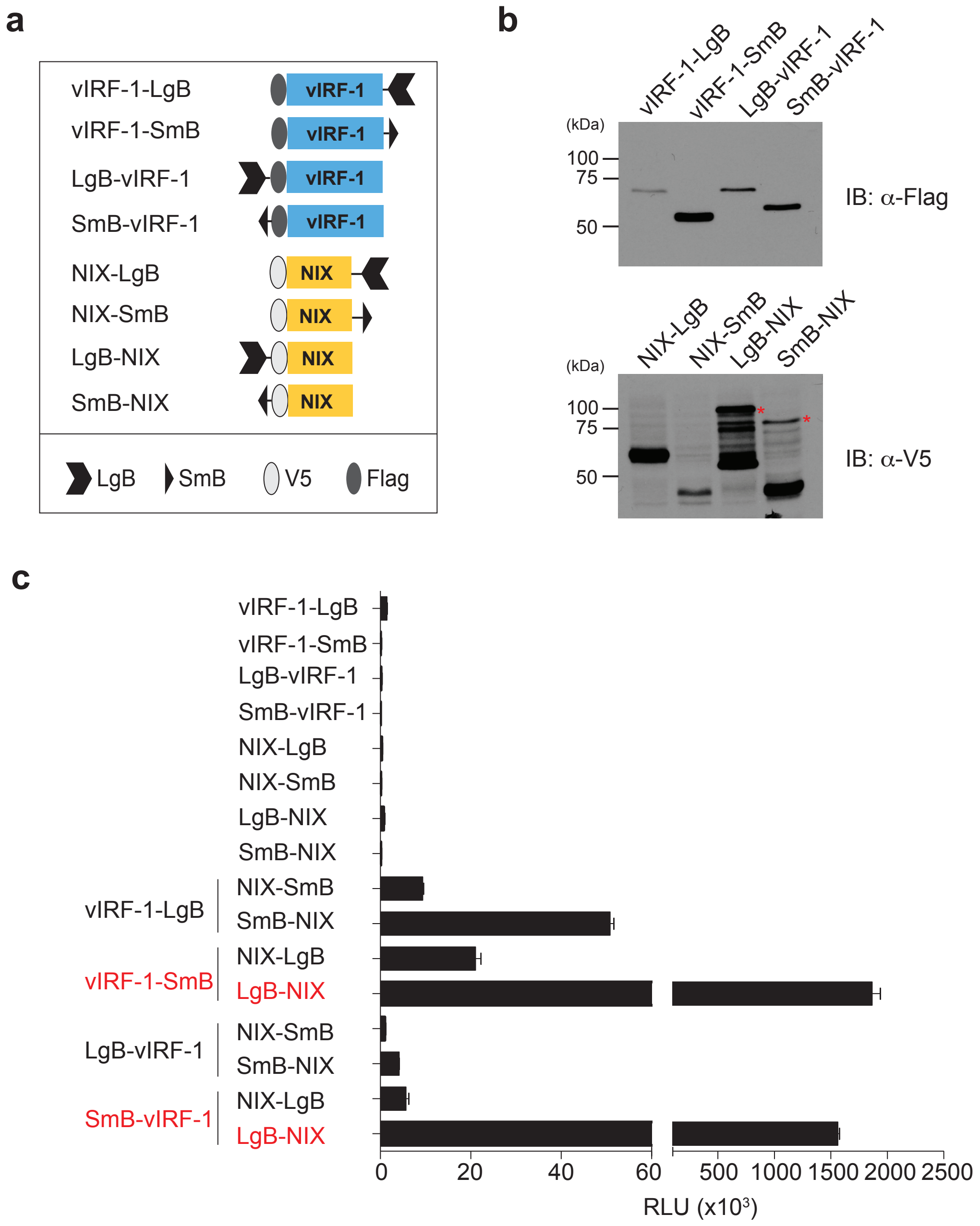

Supplementary Fig. 4

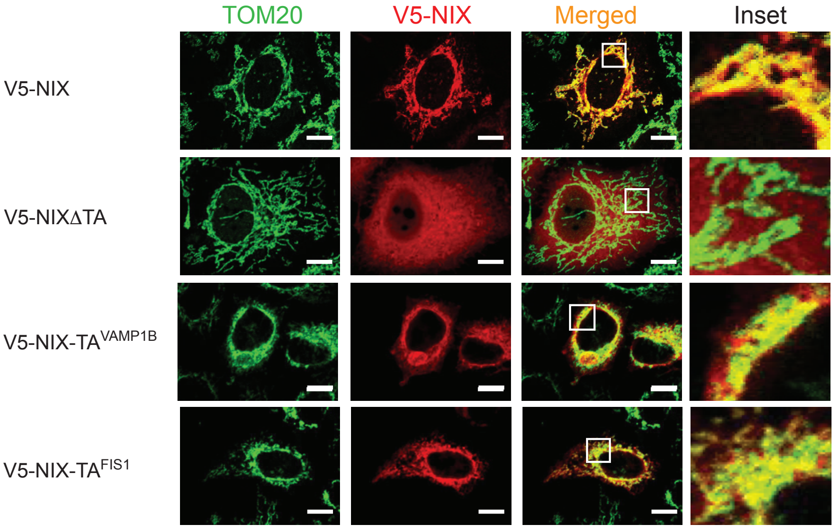

Supplementary Fig. 5

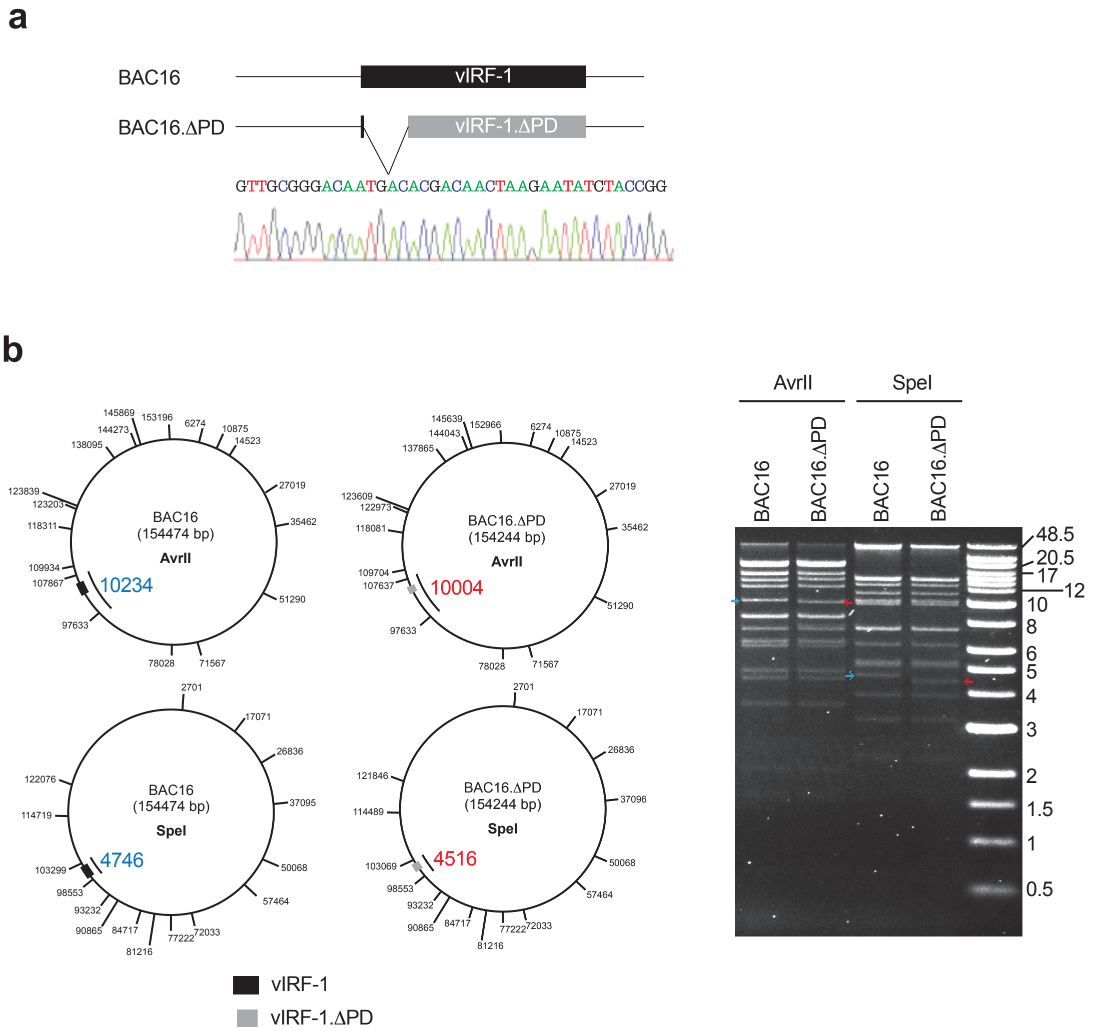

Supplementary Fig. 6

**Supplementary Table 1.** List of recombinant DNAs

| Name | Manipulation | Identifier |
|---|---|---|
| pICE_V5 | Cloning with HindIII/NotI | N/A |
| pICE_V5-NIX | Cloning with BamHI/MluI | N/A |
| pICE_V5-NIXΔLIR | Site-directed mutagenesis | N/A |
| pICE_V5-NIXΔTA | Site-directed mutagenesis | N/A |
| pICE_V5-NIX-TA$^{VAMP1B}$ | Overlap extension PCR | N/A |
| pICE_V5-NIX-TA$^{FIS1}$ | Overlap extension PCR | N/A |
| pBiT1.1N_V5-NIX | Cloning with XhoI/XbaI | N/A |
| pBiT1.1N_V5-NIXΔTA | Cloning with XhoI/XbaI | N/A |
| pBiT1.1N_V5-NIX-TA$^{VAMP1B}$ | Overlap extension PCR | N/A |
| pBiT1.1N_V5-NIX-TA$^{FIS1}$ | Overlap extension PCR | N/A |
| pBiT1.1N_Flag-vIRF-1 | Cloning with XhoI/BglII | N/A |
| pBiT1.1C_V5-NIX | Cloning with BamHI/XhoI | N/A |
| pBiT1.1C_Flag-vIRF-1 | Cloning with BglII/XhoI | N/A |
| pBiT2.1N_V5-NIX | Cloning with XhoI/XbaI | N/A |
| pBiT2.1N_Flag-vIRF-1 | Cloning with XhoI/BglII | N/A |
| pBiT2.1C_V5-NIX | Cloning with BamHI/XhoI | N/A |
| pBiT2.1C_Flag-vIRF-1 | Cloning with BglII/XhoI | N/A |
| pBiT2.1C_Flag-vIRF-1ΔPD | Cloning with BglII/XhoI | N/A |
| pcDNA3.1_vIRF-1-Flag | Lab stored | N/A |
| pcDNA3.1_vIRF-1-Flag, 18 deletion mutants | Lab stored | N/A |
| pcDNA3.1_vIRF-1 Δ2-23-Flag | Site-directed mutagenesis | N/A |
| pcDNA3.1_vIRF-1 Δ2-12-Flag | Site-directed mutagenesis | N/A |
| pcDNA3.1_vIRF-1 Δ13-23-Flag | Site-directed mutagenesis | N/A |
| pcDNA3.1_vIRF-1 Δ2-5-Flag | Site-directed mutagenesis | N/A |
| pcDNA3.1_vIRF-1 Δ6-9-Flag | Site-directed mutagenesis | N/A |
| pcDNA3.1_vIRF-1 Δ10-12-Flag | Site-directed mutagenesis | N/A |
| pcDNA3.1_vIRF-1 R6A-Flag | Site-directed mutagenesis | N/A |
| pcDNA3.1_vIRF-1 P7A-Flag | Site-directed mutagenesis | N/A |
| pcDNA3.1_vIRF-1 N8A-Flag | Site-directed mutagenesis | N/A |
| pcDNA3.1_vIRF-1 P9A-Flag | Site-directed mutagenesis | N/A |
| pcDNA3.1_vIRF-1 F10A-Flag | Site-directed mutagenesis | N/A |
| pcDNA3.1_vIRF-1 G11A-Flag | Site-directed mutagenesis | N/A |
| pGEX4T-1_NIX | Cloning with BamHI/XhoI | N/A |
| pDUET011_vIRF-1 | Cloning with BglII/XhoI | N/A |
| EZ-TET-pLKO-Puro_vIRF-1 shRNA3 (sh3-vIRF-1) | Cloning with NheI/EcoRI | N/A |
| EZ-TET-pLKO-Puro_vIRF-1 shRNA5 (sh5-vIRF-1) | Cloning with NheI/EcoRI | N/A |
| EZ-TET-pLKO-Puro_luciferase shRNA (sh-Luc) | Cloning with NheI/EcoRI | N/A |
| TET-pLKO-Neo-NIX shRNA (sh-NIX) | Cloning with AgeI/EcoRI | N/A |
| TET-pLKO-Neo-luciferase shRNA (sh-Luc) | Cloning with AgeI/EcoRI | N/A |
| BAC16.ΔPD | This study | N/A |
| NIX/BNIP3L cDNA ORF clone | Sino Biological | HG16892-U |
| pICE | Addgene | 46960 |
| pcDNA3-DRP1K38A | Addgene | 45161 |
| pDUET011 | Addgene | 17627 |
| EZ-TET-pLKO-Puro | Addgene | 85966 |
| TET-pLKO-Neo | Addgene | 21916 |

**Supplementary Table 2.** List of antibodies

| Antibody | Source | Catalog number |
|---|---|---|
| Anti-vIRF-1, rabbit | Dr. Gary Hayward | N/A |
| Anti-MTCO2, mouse | Abcam | ab110258 |
| Anti-MTCO2, rabbit | Abcam | ab79393 |
| Anti-BNIP3L/NIX, rabbit | Abcam | ab8399 |
| Anti-V5 tag-agarose, goat | Bethyl Laboratories | S190-119 |
| Anti-V5 tag, goat | Bethyl Laboratories | S190-119A |
| Anti-DYKDDDDK tag (L5) gel, rat | BioLegend | 651502 |
| Anti-BNIP3L/NIX (D4R4B), rabbit | Cell Signaling Technology | #12396 |
| Anti-LDH, rabbit | Cell Signaling Technology | #2012 |
| Anti-VDAC, rabbit | Cell Signaling Technology | #4866 |
| Anti-NBR1 (D2E6), rabbit | Cell Signaling Technology | #9891 |
| Anti-V5 tag (D3H8Q), rabbit | Cell Signaling Technology | #13202 |
| Anti-DYKDDDDK tag, rabbit | Cell Signaling Technology | #2368 |
| Anti-V5 tag (D3H8Q), rabbit | Cell Signaling Technology | #13202 |
| Anti-PARP (46D11), rabbit | Cell Signaling Technology | #9532 |
| Anti-DRP1 (D6C7), rabbit | Cell Signaling Technology | #8570 |
| Anti-HA (7C9), rat | Chromotek GMBH | 7C9-100 |
| Anti-FUNDC1, rabbit | LifeSpan BioSciences | LS-C354368 |
| Anti-p62/SQSTM1, rabbit | MBL International | PM045 |
| Anti-LC3B, rabbit | Novus Biologicals | NB100-2220 |
| Anti-β-Actin, mouse | Proteintech | 60008-1-lg |
| Anti-GST (B-14), mouse | Santa Cruz Biotechnology | sc-138 |
| Anti-TOM20 (F-10), mouse | Santa Cruz Biotechnology | sc-17764 |
| Anti-NIX (E-1), mouse | Santa Cruz Biotechnology | sc-166314 |
| Anti-Optineurin (C2), mouse | Santa Cruz Biotechnology | sc-166576 |
| Anti-Calcoco/NDP52 (F-6), mouse | Santa Cruz Biotechnology | sc-376540 |
| Anti-HSP60 (B-9), mouse | Santa Cruz Biotechnology | sc-271215 |
| Anti-ORF45 (2D4A5), mouse | Santa Cruz Biotechnology | sc-53883 |
| Anti-K8.1 (4A4), mouse | Santa Cruz Biotechnology | sc-65446 |
| Anti-mtTFA/TFAM, mouse | Santa Cruz Biotechnology | sc-376672 |
| Anti-Flag tag (M2), mouse | Sigma | F3165 |
| Anti-LAMP1, rabbit | Sino Biological | 11215-R107 |
| Anti-V5 tag, mouse | Thermo Fisher Scientific | R960-25 |
| Alexa Fluor 488-anti-mouse IgG (H+L), goat | Thermo Fisher Scientific | A11029 |
| Alexa Fluor 594-anti-rabbit IgG (H+L), goat | Thermo Fisher Scientific | A11037 |
| Alexa Fluor 647-anti-rat IgG (H+L), goat | Thermo Fisher Scientific | A21247 |
| Alexa Fluor 488-anti-mouse IgG (H+L), chicken | Thermo Fisher Scientific | A21200 |
| Alexa Fluor 594-anti-rabbit IgG (H+L), chicken | Thermo Fisher Scientific | A21442 |
| Alexa Fluor 647-anti-goat IgG (H+L), chicken | Thermo Fisher Scientific | A21469 |
| Alexa Fluor 647-anti-mouse IgG (H+L), goat | Thermo Fisher Scientific | A21236 |

**Supplementary Table 3.** List of the oligonucleotides

| Name | | Sequences (5' to 3') |
|---|---|---|
| **shRNAs** | | |
| vIRF-1 sh3 | For | ctagcggatagtatgtcaagtcaacatactagttgttgacttgacatactatcctttttg |
| | Rev | aattcaaaaaggatagtatgtcaagtcaacaactagtatgttgacttgacatactatccg |
| vIRF-1 sh5 | For | ctagcggcattctgctgactagctcttactagtagagctagtcagcagaatgcctttttg |
| | Rev | aattcaaaaaggcattctgctgactagctctactagtaagagctagtcagcagaatgccg |
| Luc shRNA (I) | For | ctagccctaaggttaagtcgccctcgtactagtcgagggcgacttaaccttaggtttttg |
| | Rev | aattcaaaaacctaaggttaagtcgccctcgactagtacgagggcgacttaaccttaggg |
| NIX shRNA | For | ctagccagtcagaagaagaagttgtatactagttacaacttcttcttctgactgtttttg |
| | Rev | aattcaaaaacagtcagaagaagaagttgtaactagtatacaacttcttcttctgactgg |
| Luc shRNA (II) | For | ccggtcctaaggttaagtcgccctcgtactagtcgagggcgacttaaccttaggtttttg |
| | Rev | aattcaaaaacctaaggttaagtcgccctcgactagtacgagggcgacttaaccttagga |
| **RT-qPCR primers** | | |
| NIX | For | ctacccatgaacagcagcaa |
| | Rev | atctgcccatcttcttgtgg |
| TFAM | For | ccgaggtggttttcatctgt |
| | Rev | tccgccctataagcatcttg |
| 18S RNA | For | gtaacccgttgaaccccatt |
| | Rev | ccatccaatcggtagtagcg |
| LANA | For | tacggttggcgaagtcacatc |
| | Rev | cctcgcagcagactacacctccac |
| **Cloning** | | |
| V5 tag pICE | For | agcttgccaccggtaagcctatccctaaccctctcctcggtctcgattctacgtgc |
| | Rev | ggccgcacgtagaatcgagaccgaggagagggttagggataggcttaccggtggca |
| NIX-TA VAMP1B | For | tctccgcagaatttctgaagatgatgatcatgctgggagccatctgtgccatcatc |
| | Rev | gagggtgtgctcagtcgctttacaataactaccacgatgatggcacagatggctcc |
| NIX-TA FIS1 | For | tctccgcagaatttctgaagctcgtgggcatggccatcgtgggaggcatggccctgggtg |
| | Rev | gagggtgtgctcagtcgcttgatgagtccggccagtcccgccacacccagggccatgcct |
| **Mutagenesis** | | |
| BAC16 ΔPD | For | tggacattgcggcgcgagctagtctggttgcgggacaatgacacgacaactaagaa tatcaggatgacgacgataagtaggg |
| | Rev | tcccctcactggcaccggtagatattcttagttgtcgtgtcattgtcccgcaacca gactcaaccaattaaccaattctgattag |
| **PCR primers for mitochondrial FISH probes** | | |
| mtDNA (1) | For | TCCATGCATTTGGTATTTTCGTC |
| | Rev | CGAAGGGTTGTAGTAGCCCGTAG |
| mtDNA (2) | For | ACCTTCAAATTCCTCCCTGTACG |
| | Rev | TGATGGCCCCTAAGATAGAGGAG |
| mtDNA (3) | For | GGCAACCTTCTAGGTAACGACCA |
| | Rev | GAGGAGCGTTATGGAGTGGAAGT |
| mtDNA (4) | For | GTAAGCCTCTACCTGCACGACAA |
| | Rev | GTGGATGCGACAATGGATTTTAC |
| mtDNA (5) | For | GCTCACTCACCCACCACATTAAC |
| | Rev | TGAAGGGCAAGATGAAGTGAAAG |
| mtDNA (6) | For | TGAGGGGCCACAGTAATTACAAA |
| | Rev | GAGTGGGAGGGGAAAATAATGTG |
